# Polymeric mechanism of enhancer-promoter cooperativity in transcriptional bursting

**DOI:** 10.64898/2026.04.29.721797

**Authors:** Tetsuya Yamamoto, Koji Kawasaki, Takashi Fukaya

**Affiliations:** Institute for Chemical Reaction Design and Discovery, Hokkaido University, Sapporo 010-0021, Japan; Laboratory of Transcription Dynamics, Research Center for Biological Visualization, Institute for Quantitative Biosciences, The University of Tokyo, Bunkyo-ku, Tokyo 113-0032, Japan; Department of Life Sciences, Graduate School of Arts and Sciences, The University of Tokyo, Bunkyo-ku, Tokyo 113-0032, Japan

## Abstract

Emerging evidence suggest that gene expression is controlled though the modulation of transcriptional bursting across species. However, the underlying regulatory mechanisms remain largely uncertain. Recent live-imaging studies have reported that transcription factors (TFs) form a cluster at enhancers just prior to gene activation, thereby locally concentrating the active transcription machinery. This process is thought to be mediated by multivalent interaction between Mediator recruited by TFs. Conversely, transcription itself is also suggested that to influence the assembly of the TF clustering, implicating the presence of a feedback mechanism. To understand the theoretical framework underlying the interplay between TF clustering and transcription, we here develop a polymer micelle model of transcriptional bursting. With this model, multiple RNA polymerase II (Pol II) molecules loaded to the promoter, together with enhancer-bound Mediator, assemble a micelle-like structure due to their connectivity via DNA when the chromatin fiber connecting enhancer and promoter adopts closed conformation, analogous to polysoap micelle. This assembly further recruits freely diffusing Pol II and Mediator in the nucleoplasm even at low concentration up to the optimal size. Our theoretical framework enables quantitative prediction of how dynamic transitions of enhancer-promoter conformation and the stability of the micelle impact the kinetics of transcriptional bursting.

## Introduction

Spatial and temporal specificity of gene transcription is dynamically controlled by distal enhancers located at genomic distances away from their target genes^1-6^. Disruption of these regulatory interplay has been implicated in various genetic diseases^7-9^. Enhancers serve as binding sites of sequence-specific transcriptional factors (TFs), which subsequently recruit cofactors such as Mediator^3-5,10^. Classically, transcription has been thought to be activated through physical association between distal enhancers and target promoters via looping out intervening DNA sequences^10-13^. However, this classical model has been challenged by recent studies showing that transcription activity is not directly proportional to the enhancer-promoter contact frequency^14^ and that the enhancer-promoter 3D distances do not necessarily correlate with transcription states^15-17^.

Kinetically, transcription occurs in intermittent bursts, in which multiple Pol II molecules are recruited to promoters and initiate productive elongation in a coordinated manner during the ON state followed by a transcriptionally quiescent OFF state^18-23^. We have experimentally shown that TFs assemble into cluster just prior to the onset of transcriptional bursting and rapidly disassemble before entering the refractory OFF state^18^. The number of TF molecules enriched within a cluster is shown to be larger than the number of their binding sites at the enhancer, raising the possibility that the major driving force behind cluster assembly is multivalent protein-protein interaction, including TFs and Mediator^24,25^.

Interestingly, TF clusters seeded at enhancer regions have been shown to grow during the ramp-up phase of transcriptional bursting and to be sensitive to linear genomic distance between enhancers and promoters^18^, implicating that the assembly of TF clusters is controlled by cooperative actions between enhancers and promoters. Similar implication was obtained by recent Micro-C experiments^26^. However, molecular mechanism underlying this feedback process between TF clustering and transcription remains mostly unclear. To address this question, theoretical predictions of the interplay between TF clustering and transcriptional bursting have strong potential to elucidate the kinetic control of gene transcription by distal enhancers.

A key molecular foundation of transcriptional bursting is that multiple Pol II molecules are loaded onto the promoter and initiate productive elongation in a coordinated manner. Structural studies provided direct molecular evidence that Mediator associates with the C-terminal domain (CTD) of Pol II to form a complex^27,28^. Thus, it is reasonable to assume Pol II-Mediator complexes are locally multimerized at active gene loci through association with a single chromatin fiber. These multimerized Pol II-Mediator complexes can be further stabilized though multivalent protein-protein interactions involving the intrinsically disordered regions of Mediator subunits^24,25^. This situation is analogous to a polysoap, in which hydrophilic heads of surfactant units are connected by a hydrophilic chain^29^. In contrast to regular surfactants (Fig. 1**a**), polysoaps readily assemble into micelles regardless of their concentration, because the hydrophobic tails of the surfactant units are already in close spatial proximity^29,30^ (Fig. 1**b**). These micelles can grow by incorporating regular surfactants mixed with polysoaps, even at concentrations lower than the critical micelle concentration (CMC), up to an optimal size^26^ (Fig. 1**c**). Likewise, we hypothesize that multiple Pol II-Mediator complexes recruited to promoter and enhancer regions assemble into micelle-like structure due to their molecular similarity to polysoaps. The resulting micelle may further grow by incorporating freely diffusing Pol II and Mediator in the nucleoplasm, ultimately giving rise to the activator clusters observed in recent live-imaging studies^18^.

**Figure 1.**
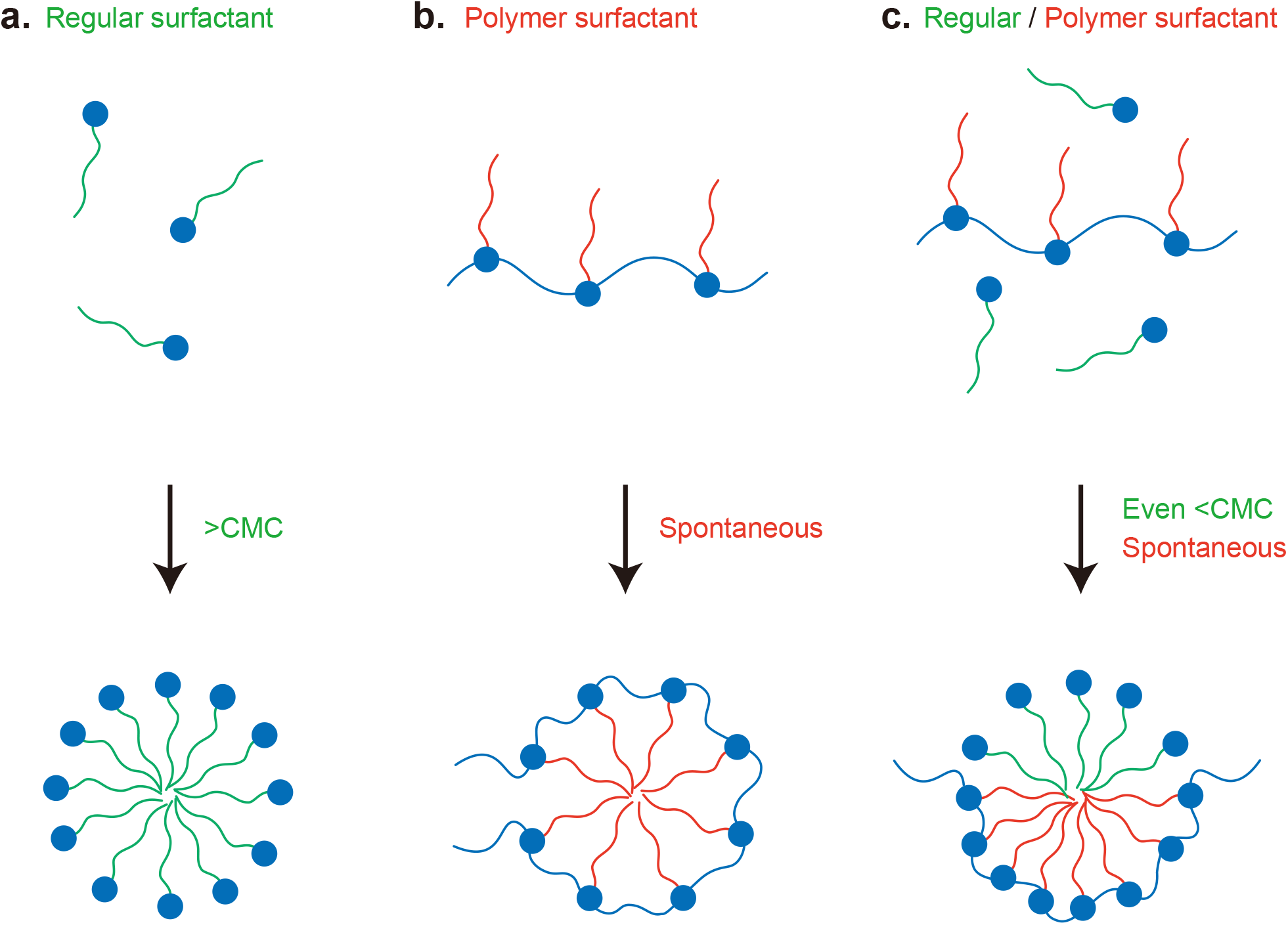
Polysoap micellization. **a**. Regular surfactant is compounds synthesized by covalently bonding hydrophobic chains to a hydrophilic head. Regular surfactants in an aqueous solution assemble micelles at the concentrations higher than critical micelle concentration (CMC). **b**. Polysoap is a polymer synthesized by connecting the hydrophilic heads of surfactant units by a hydrophilic chain. Polysoaps readily assemble micelles in an aqueous solution regardless of concentration because of the connectivity. **c**. Regular surfactants mixed with polysoaps are incorporated into micelles of polysoaps at the concentrations lower than CMC.

Based on this biophysical perspective, we develop a theoretical framework for transcriptional bursting by integrating the kinetics of biochemical processes involved in transcription into the soft matter physics of polysoap micellization^30^. Our theory identifies the activator clusters as micelles of Pol II-Mediator complexes assembled onto chromatin. We further suggest that the ON and OFF states of transcriptional bursting are functionally linked to the assembled and disassembled states of the micelle of Pol II-Mediator complexes (Fig. 2**a**). Our theory enables us to quantitatively predict the kinetic parameters of transcriptional bursting based on the stability of the micelle and the number of complexes bound at promoter and enhancer regions. Importantly, key biological properties of enhancers including nonlinear regulation of transcription through interaction with promoters^14^ can be recapitulated within this framework. We believe that our theory will serve as a key foundation for designing future experiments to elucidate the biophysical principles underlying transcriptional bursting and 3D genome organization.

**Figure 2.**
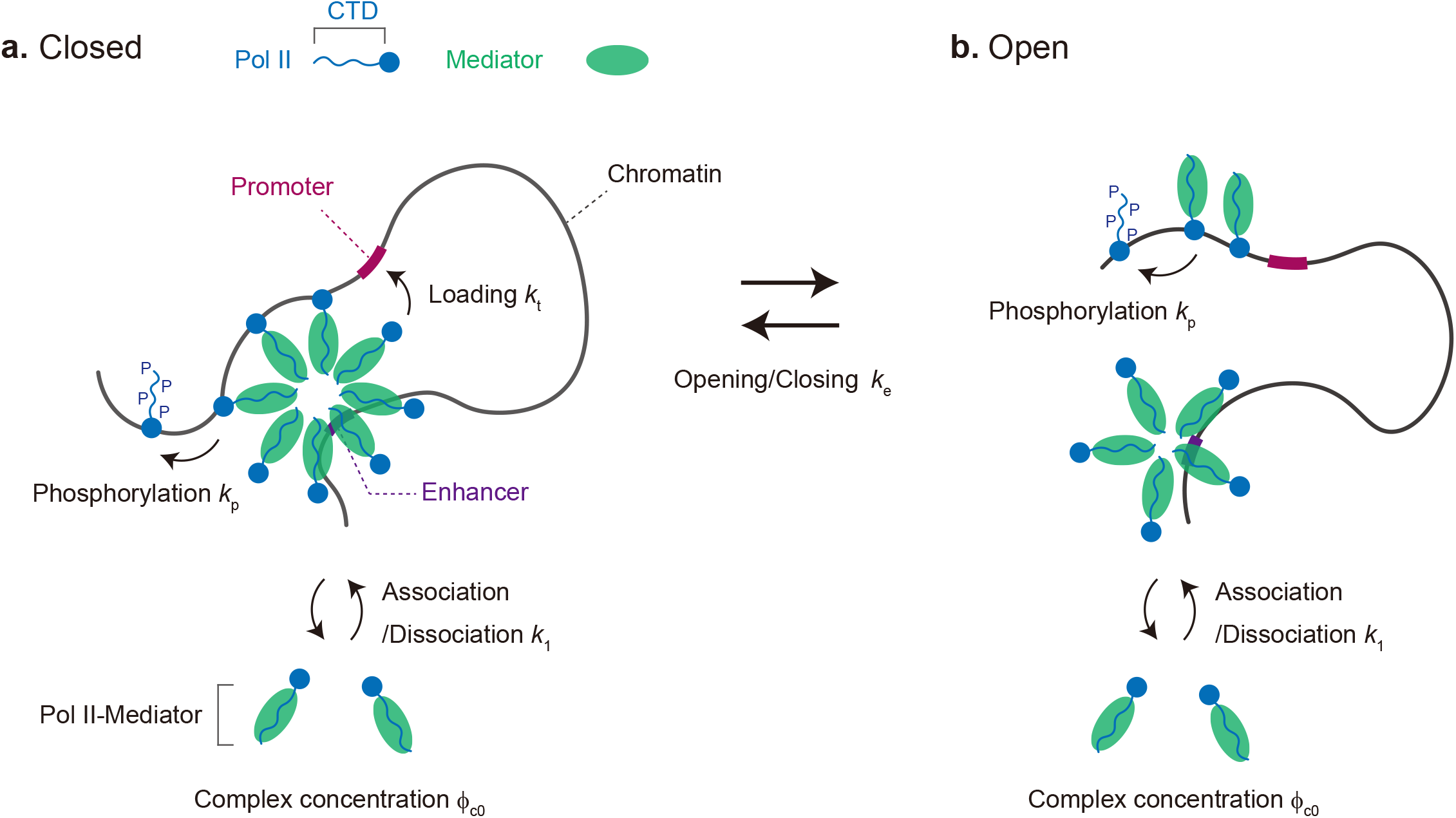
Polymer model of transcriptional burst. We consider a system of the DNA region (black) between enhancer and promoter (thick brown lines). The genomic distance between these units is measured by the number *N*_d_ of Kuhn units. Complexes of Mediator (green) and Pol II (blue) bind to the binding sites in the enhancer and assemble a micelle there. The DNA region forms the closed (**a**) or open (**b**) conformation. Complexes in the micelle are loaded to the promoter for transcription only at the closed conformation. The state of the system is described by the conformational state of the DNA region, the number *m* of complexes in the micelle, and the number *n* of complexes bound to the promoter.

## Results

### Polymer micelle model of transcriptional bursting

We develop a theoretical model of transcriptional bursting that incorporates the loading of Pol II-Mediator complexes to the promoter, phosphorylation of the Pol II CTD, transitions between open and closed conformations of chromatin fiber connecting enhancer and promoter, and the assembly and disassembly of micelle of Pol II-Mediator complexes (Fig. 2). Our theory represents the state of the system by the total number *m* of Pol II-Mediator complexes in the micelle, the number *n* of complexes loaded onto the promoter for transcription, and the conformational state of the chromatin fiber. In our model, state transitions are described by using a master equation, which accounts for fluctuations in both *m* and *n* (Eqs. 4 and 5).

In this model, the enhancer is treated as binding sites of Pol II-Mediator complexes, as suggested by recent single-molecule imaging studies^31,32^. Based on our previous observation that Mediator forms clusters at enhancer regions^18^, we assume that the micelle assembles at the enhancer rather than the promoter. To describe the kinetics of micellar assembly and disassembly, we extend the pioneering work by Aniansson and Wall^33^, which assumes stepwise association and dissociation of complexes. The association rate is proportional to the volume fraction *ϕ*_c0_ of complexes in the nucleoplasm and the number of unoccupied binding sites at the enhancer, until all sites are fully occupied (first terms of eqs. (6) and (7)). This treatment reflects the correlation arising from the connectivity of binding sites within the enhancer^34-36^. The dissociation rate depends on the free energy *F*_*m,n*_ of the micelle (second terms of eqs. (6) and (7)), which accounts for multivalent Mediator-Mediator interactions and excluded volume interactions between Pol II molecules (eq. (1)).

A stretch of chromatin fiber connecting the promoter and the enhancer can adopt either a closed (Fig. 2**a**) or open (Fig. 2**b**) conformation. We derive the transition rate from the open to closed conformation by treating chromatin as an ideal chain^34,35,37^, analogous to the treatments of the corresponding transition of telechelic polymers^38^ (eq. (11)). In our model, the linear enhancer-promoter genomic distance *N*_d_, measured in units of Kuhn segments, affects the output only through the closing rate (eq. (10)).

In the closed conformation of chromatin fiber connecting the enhancer and promoter elements, complexes in the micelle are loaded onto the promoter due to their spatial proximity (Fig. 2**a**). The loading rate is proportional to the number *m* − *n* of complexes that are in the micelle and have not been loaded to the promoter (third and fourth terms of eq. (4)). The complexes already loaded onto the promoter are connected via chromatin, analogous to polysoap; thus, they can be incorporated into the micelle without incurring a translational entropy cost^30^. The CTDs of promoter-loaded Pol II molecules are subsequently phosphorylated, leading to dissociation of Mediator from the complexes^39^ (Fig. 2**a**). Concurrently with Mediator dissociation, phosphorylated form of Pol II enters into productive elongation and becomes less adhesive, thereby dissociating from the micelle (fifth and sixth term of eq. (4) and third and fourth terms of eq. (5)). The number *n* of loaded complexes is determined by the balance of the loading and dissociation rates.

The opening rate of the enhancer and promoter elements is calculated based on the free energy cost required to dissociate *n* complexes from the micelle (eighth term of eq. (4) and sixth term of eq. (5)). In the open conformation, the promoter is not in spatial proximity to the micelle assembled at the enhancer; complexes are no longer loaded onto the promoter (Fig. 2**b**). Within this framework, freely diffusing Pol II molecules are assumed not to be promiscuously loaded onto the promoter independently of the enhancer, based on past observations that minimal core promoters rarely activate transcription in the absence of linked activator binding sites or enhancers^31,32^. Accordingly, we neglect enhancer-independent transcription, in which freely diffusing Pol II molecules might otherwise be loaded onto the promoter. Because the promoter-bound complexes are located far from the micelle, the micelle has a higher probability of disassembling in the open conformational state.

### Linking theory and experiments

If the loading of Pol II molecules onto the promoter and the subsequent phosphorylation reactions are halted with a fixed number *n* of loaded complexes, the association and dissociation rates of the complexes satisfy the detailed balance, and our theory reduces to the kinetics of micellar formation of polymer surfactants (first and second terms of eqs. (4) and (5)). According to the second law of thermodynamics, at thermodynamic equilibrium, the most stable state corresponds to the minimum of the free energy difference ℱ_*m,n*_ to assemble a micelle from dispersed individual complexes (see Supplementary equation (21)). The free energy difference ℱ_*m,n*_ typically exhibits a maximum at *m* = *m*^∗^, which acts as a free energy barrier separating two minima at *m* = *m*_1_ (< *m*^∗^) and *m* = *m*_2_ (> *m*^∗^) (Fig. 3**a**)^38,40^. The regions *m* ≤ *m*^∗^ and *m* > *m*^∗^ correspond to the disassembled and assembled states of the micelle, respectively.

**Figure 3.**
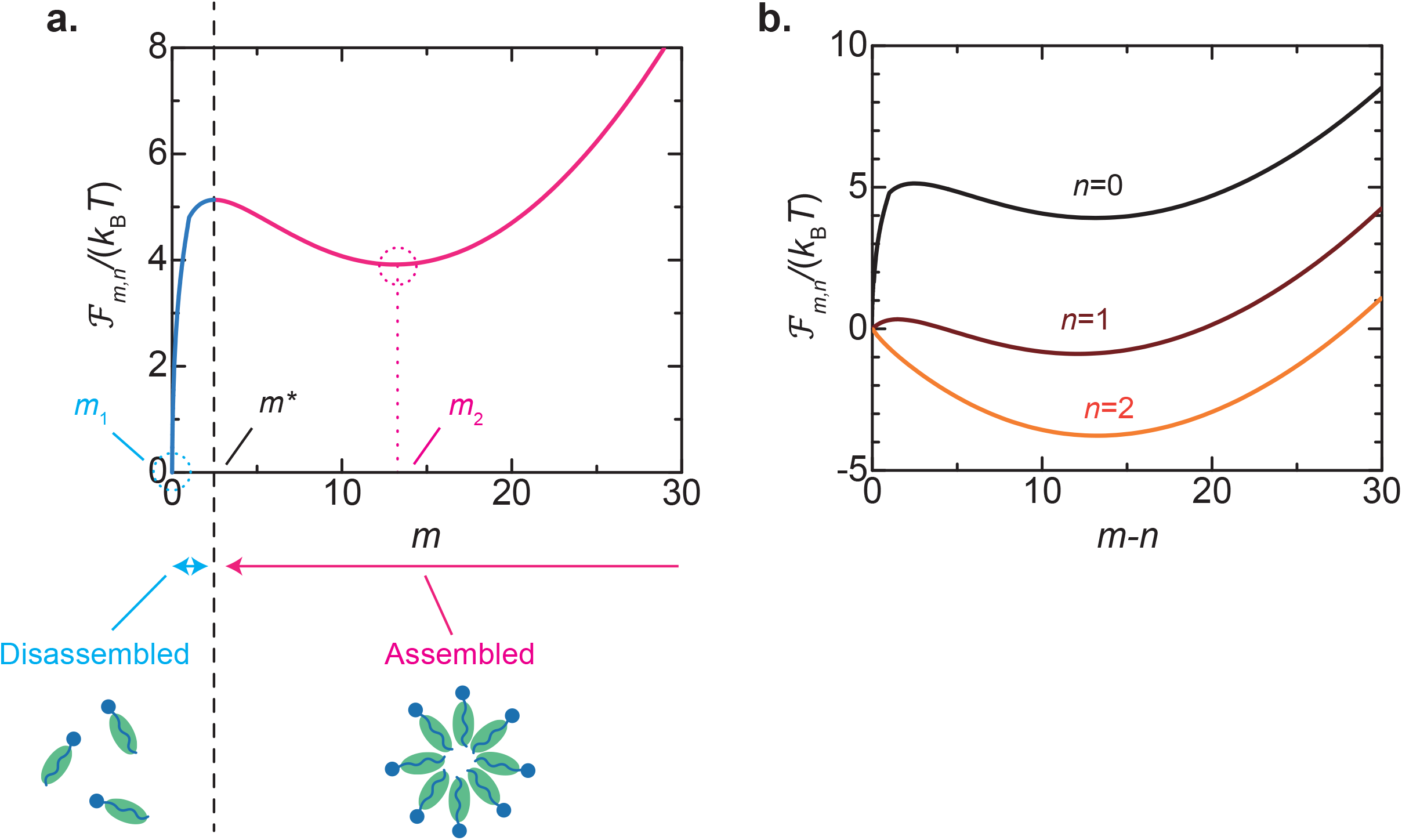
Free energy difference ℱ_*mm*_. The free energy difference ℱ_*mn*_ (rescaled by the thermal energy *k*_B_*T*) is shown as a function of the number *m* of Pol II-Mediator complexes in the micelle for *n* = 0 (**a** and black line in **b**), 1 (brown line in **b**), and 2 (orange line in **b**) in the closed conformation. **a**. The free energy difference typically has two local minima, *m*_1_ and *m*_2_, and one maximum *m*^∗^. The regions, *m* ≤ *m*^∗^ and *m* > *m*^∗^, are identified as the disassembled and assembled states of micelle, respectively. **b**. The assembled state becomes more stable, while the disassembled state becomes more unstable, as the number *n* of complexes loaded to the promoter increases. The values of parameters used for the calculations are summarized in Table 2.

In the closed conformation of the chromatin fiber between enhancer and promoter elements, the assembled state becomes more stable, whereas the disassembled state becomes less stable as the number *n* of complexes loaded onto the promoter increases (Fig. 3**b**). Based on our previous live-imaging study showing that transcriptional bursting coincides with the dynamic assembly of activator cluster at the enhancer^16^, we identify the assembled and disassembled states of the micelle as the ON and OFF states of transcriptional bursting (Fig. 4**a**). Importantly, the ON and OFF states are, by definition, independent of the closed and open conformational states of the chromatin fiber between the enhancer and promoter elements. Thus, the active gene loci can, in principle, adopt either the closed or open conformation within each micellar state.

**Figure 4.**
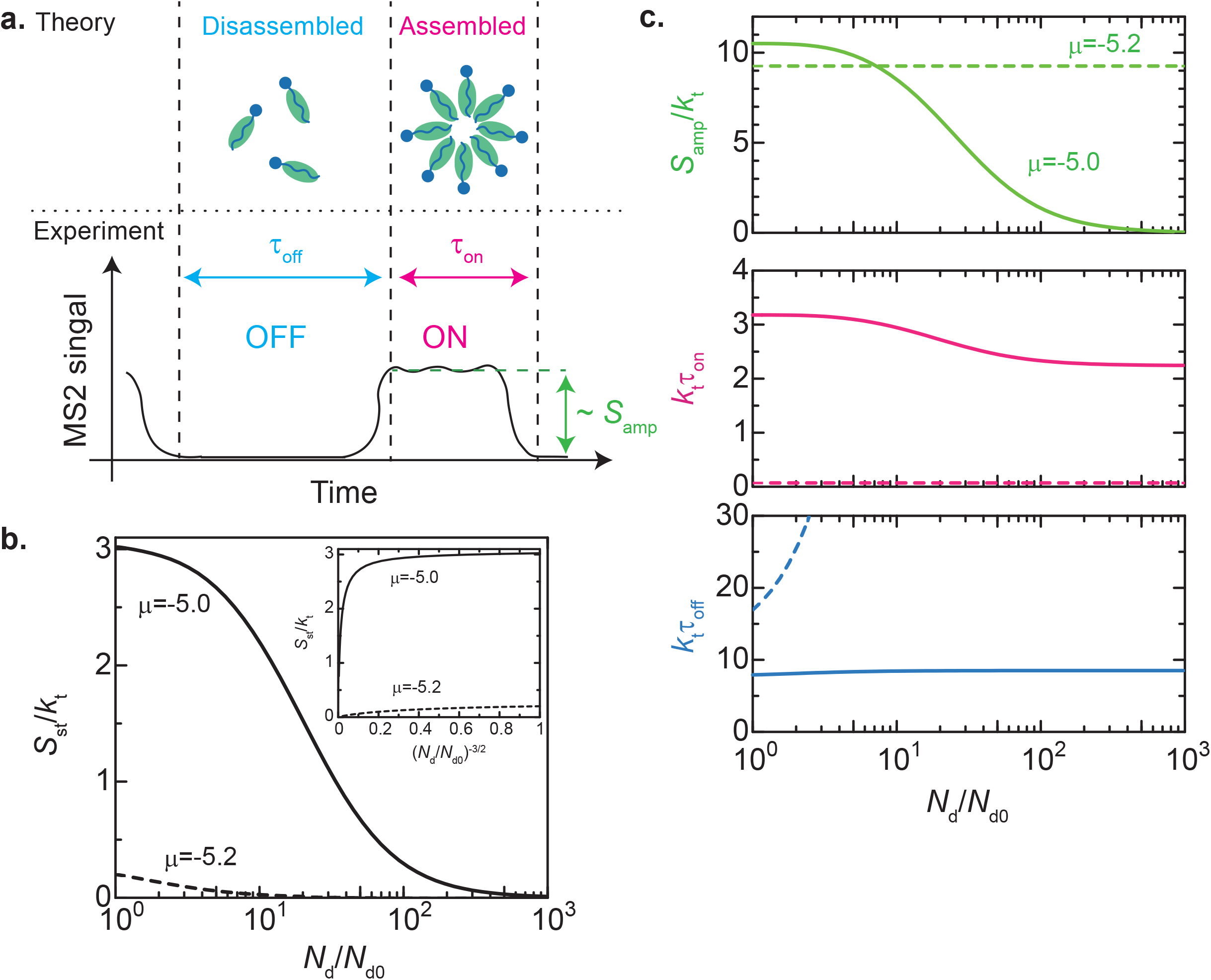
Transcription rate vs linear enhancer-promoter genomic distance *N*_d_. **a**. The polymer model predicts the burst size *S*_amp_, ON duration *τ*_on_, and OFF duration *τ*_off_, identifying the ON and OFF states of transcriptional burst as the assembled and disassembled states of micelle, respectively. **b**. The loading rate *S*_st_ of complexes in the steady state (rescaled by loading rate constant *k*_t_) is shown as a function of the linear enhancer-promoter genomic distance *N*_d_ (rescaled by *N*_d0_) for *μ* = −5.0 (solid line) and −5.2 (broken line). (Inset) The linear enhancer-promoter distance *N*_d_ is converted to a quantity 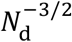, proportional to the contact frequency. **c**. The burst size *S*_amp_ (light green), ON duration *τ*_on_ (magenta), and OFF duration *τ*_off_ (cyan) (rescaled by the rate constant *k*_t_) are shown as functions of the linear genomic distance *N*_d_ (rescaled by *N*_d0_) for *μ* = −5.0 (solid line) and −5.2 (broken line). The value of parameters used for the calculations are summarized in Table 2.

The free energy difference ℱ_*mn*_ exhibits a single minimum *m* = *m*_1_ (≈ 0) when the number *n* of complexes loaded onto the promoter is below a threshold *n*_sp1_ and the complex volume fraction is smaller than a threshold, 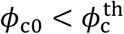 (Supplementary Figure 1). In contrast, it exhibits a single minimum *m* = *m*_2_ (≫ 1) when *n* is larger than another threshold *n*_sp2_ in the closed conformation of the chromatin fiber connecting the promoter and enhancer elements (Fig. 3**b**; orange line). The loading of complexes up to *n* = *n*_sp2_ and the phosphorylation of complexes at *n* = *n*_sp1_ provide additional pathways for regulating the assembly and disassembly dynamics of the micelle. In the open conformation of the chromatin fiber connecting the enhancer and promoter elements, the complexes loaded onto the promoter do not contribute to the stability of the micelle assembled at the enhancer. When 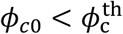, the micelle disassembles upon loss of promoter-loaded complexes.

To build our theoretical framework, we introduce many parameters to account for micellization, the biochemical reactions involved in transcriptional activation, and enhancer function (Table 1). It should be emphasized that most of these parameters are physical and chemical quantities that can, in principle, be experimentally characterized, rather than fitting parameters. Moreover, the predictions of the model depend only on ratios of time, length, and energy scales, thereby reducing the number of independent parameters (Table 2). Thus, our theory provides a generalizable and testable framework that quantitatively predict gene expression output.

**Table 1.** Native parameters involved in our theory.

| Symbol | Meaning | Estimate | Eq | Ref |
| --- | --- | --- | --- | --- |
| $\gamma$ | Surface energy per unit area | 16 $\mu\text{N/m}$ | 1 | |
| $K$ | Pol II-Pol II interaction | $2 \times 10^{-35} \text{ J}\cdot\text{m}^2$ | 1 | |
| $\chi_m$ | Med-Med interaction (per $k_B T$) | $< 7$ | 1 | |
| $r_1$ | Pol II CTD-Med size | 16 nm | 2 | 41 |
| $k_t$ | Rate constant: Complex loading | $0.2 \text{ s}^{-1}$ | 4,5 | 23 |
| $k_p$ | Rate constant: Complex phosphorylation | - | 4,5 | |
| $k_e$ | Rate constant: Conformational transition | - | 4,5 | |
| $k_1$ | Rate constant: Complex association | $5 \times 10^3 \text{ s}^{-1}$ | 6-9 | |
| $\phi_{c0}$ | Complex volume fraction | 0.001 | 6-9 | |
| $\epsilon_e$ | Enhancer binding energy (per $k_B T$) | $\sim -1$ | 4,5 | |
| $n_e$ | Number of enhancer binding sites | Variable | 6,7 | |
| $b$ | Chromatin unit length | 50 nm | 11 | 42 |
| $N_d$ | Enhancer-promoter genomic length | Variable | 11 | |
| $\Delta$ | Cutoff distance | $\sim 50 \text{ nm}$ | 10 | |

**Table 2.**
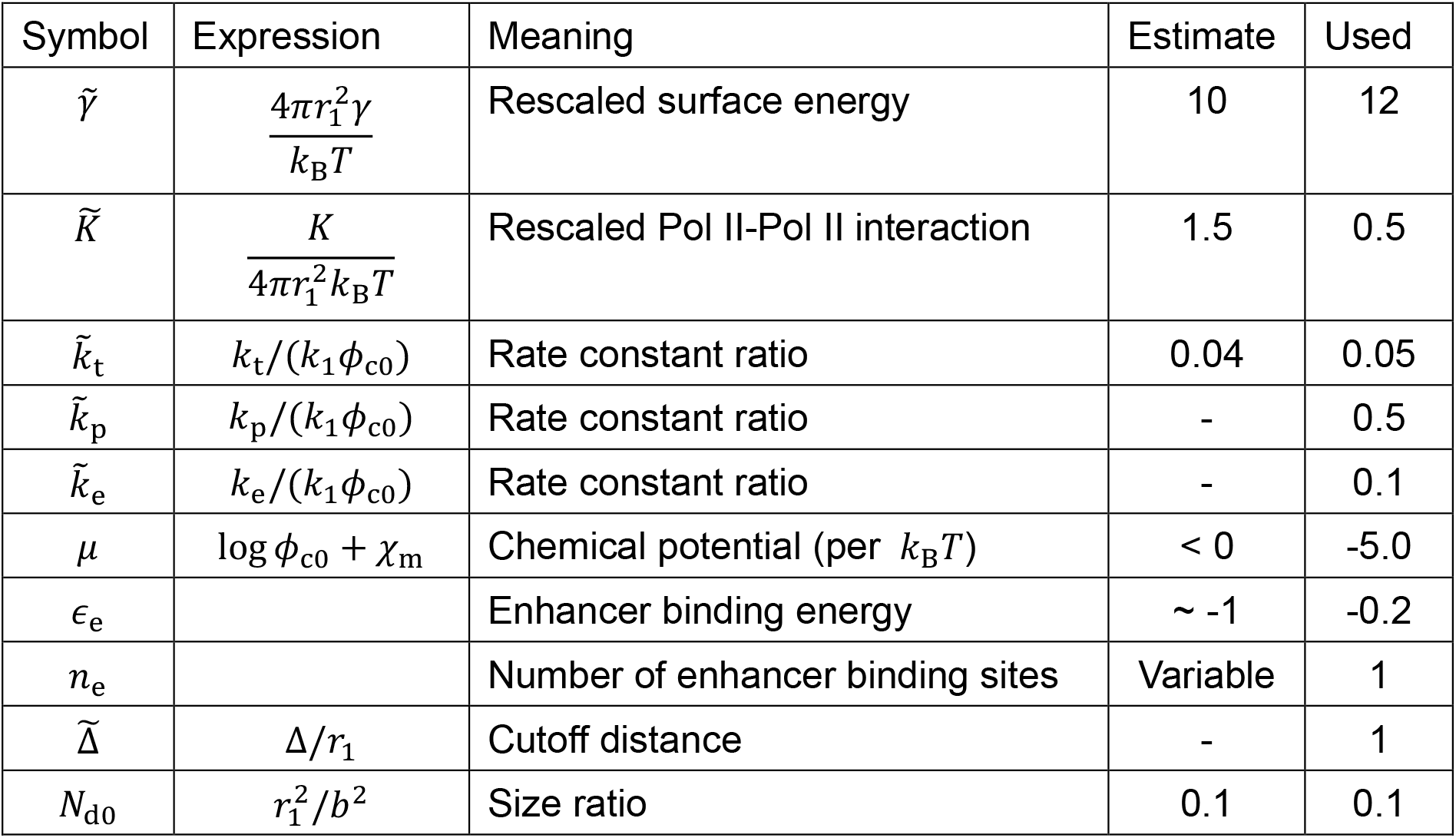
Ratios of time, length, and energy scales.

The surface energy per unit area is estimated by using the thermal energy, 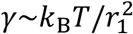 (see refs. 40 and 43). The elastic constant *K* that accounts for the Pol II-Pol II interaction is estimated by using 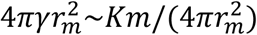 by estimating the number *m* of complexes in the micelle as ∼20. The diffusion constant of complexes is estimated by using the Einstein-Stokes relationship^37^, *D*_1_∼*k*_*B*_*T*/(6*πη*_s_*r*_1_), where *η*_s_ (∼ 0.01 Pa ⋅ s) is the viscosity of nucleoplasm (estimated from the viscosity of egg extract^44^). The loading rate is estimated as the inverse of the relaxation time of the complex, 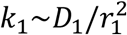. The concentration of complexes in the solution is estimated as 100 nM. One Kuhn unit of chromatin corresponds to 1.65 kbps^42^.

The size ratio *N*_d0_ (∼ 0.1) corresponds to 0.165 kbps.

### The stability of micelle governs kinetic parameters of transcriptional bursting

We first analyze the steady-state transcription rate *S*_st_, which corresponds to the average over the assembled (ON) and disassembled (OFF) states (Supplementary equation (88)). Our theory predicts that *S*_st_ decreases non-linearly with increasing linear enhancer-promoter genomic distance *N*_d_ (Fig. 4**b**), consistent with previous studies^14,45^. For ideal chains, the enhancer-promoter contact frequency scales as 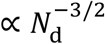. The transcription rate *S*_st_ is a sigmoidal function of the contact frequency, further supporting the non-linearity function of enhancers in controlling gene expression, as suggested by recent experiments^14,45^, at least qualitatively (Fig. 4**b**; inset).

Our theory can also predict kinetic parameters of transcriptional bursting, such as burst size *S*_amp_ and durations, *τ*_on_ and *τ*_off_ (Supplementary equation (83) – (85)). For the case of 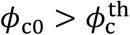, the burst size *S*_amp_ and the duration *τ*_on_ of the ON state decrease with increasing the linear enhancer-promoter genomic distance *N*_d_ (Fig. 4**c**; green and magenta solid lines). The duration *τ*_off_ of the OFF state, by contrast, decreases with increasing *N*_d_ (Fig. 4**c**; blue solid line). This behavior reflects the fact that, in this regime, the assembled state of the micelle is metastable in the open conformation of the chromatin fiber between the enhancer and promoter elements but the re-establishment of the closed conformation can rescue the micelle from disassembly. In contrast, for the case of 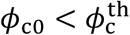, the burst size and the duration of the ON state remain constant (Fig. 4**c**; green and cyan dot lines). The duration of the OFF state *τ*_off_ increases with increasing *N*_d_ (Fig. 4**c;** blue dot line). This reflects the fact that, in this regime, the assembled state of the micelle is unstable in the open conformation of the chromatin fiber between the enhancer and promoter elements, and the micelle cannot be rescued from disassembly once the enhancer and promoter separate. Thus, our theory predicts that the kinetic parameters of transcriptional bursting are highly sensitive to the factors that influence in micelle stability, such as the concentration *ϕ*_c0_ of Pol II-Mediator complexes.

### Enhancer-promoter cooperativity is tuned by the phosphorylation of Pol II CTD

Our theory assumes that enhancer-promoter cooperativity results from the connectivity of Pol II-Mediator complexes loaded onto the promoter. The number *n* of the loaded complexes increases with decreasing the rate *k*_p_, at which Mediator dissociates from Pol II upon phosphorylation of CTD^39^ (Supplementary Figure 2**a**). Consistent with this, the mean number 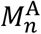 of complexes comprising the micelle increases with decreasing *k*_p_, supporting a model in which enhancer-promoter cooperativity is mediated through the connectivity of Pol II-Mediator complexes loaded onto the promoter (Supplementary Figure 2**b**).

Our theory predicts that the burst size *S*_amp_ and the duration *τ*_on_ of the ON state increase as the phosphorylation rate *k*_p_ decreases (Fig. 5; green and magenta solid lines). Importantly, the dependence of *S*_amp_ on the linear enhancer-promoter genomic distance *N*_d_ becomes less evident as *k*_p_ decreases (Fig. 5; green solid and dot lines). This is because the closed conformation of the chromatin fiber between the enhancer and promoter elements is stabilized by the complexes bound to the promoter. The duration *τ*_off_ of the OFF state decreases with decreasing *k*_p_ (Fig. 5; blue solid and broken lines). Overall, our results are consistent with the idea that the kinetic parameters of transcriptional bursting are controlled by the rate of Pol II CTD phosphorylation and the number *n* of complexes bound to the promoter.

**Figure 5.**
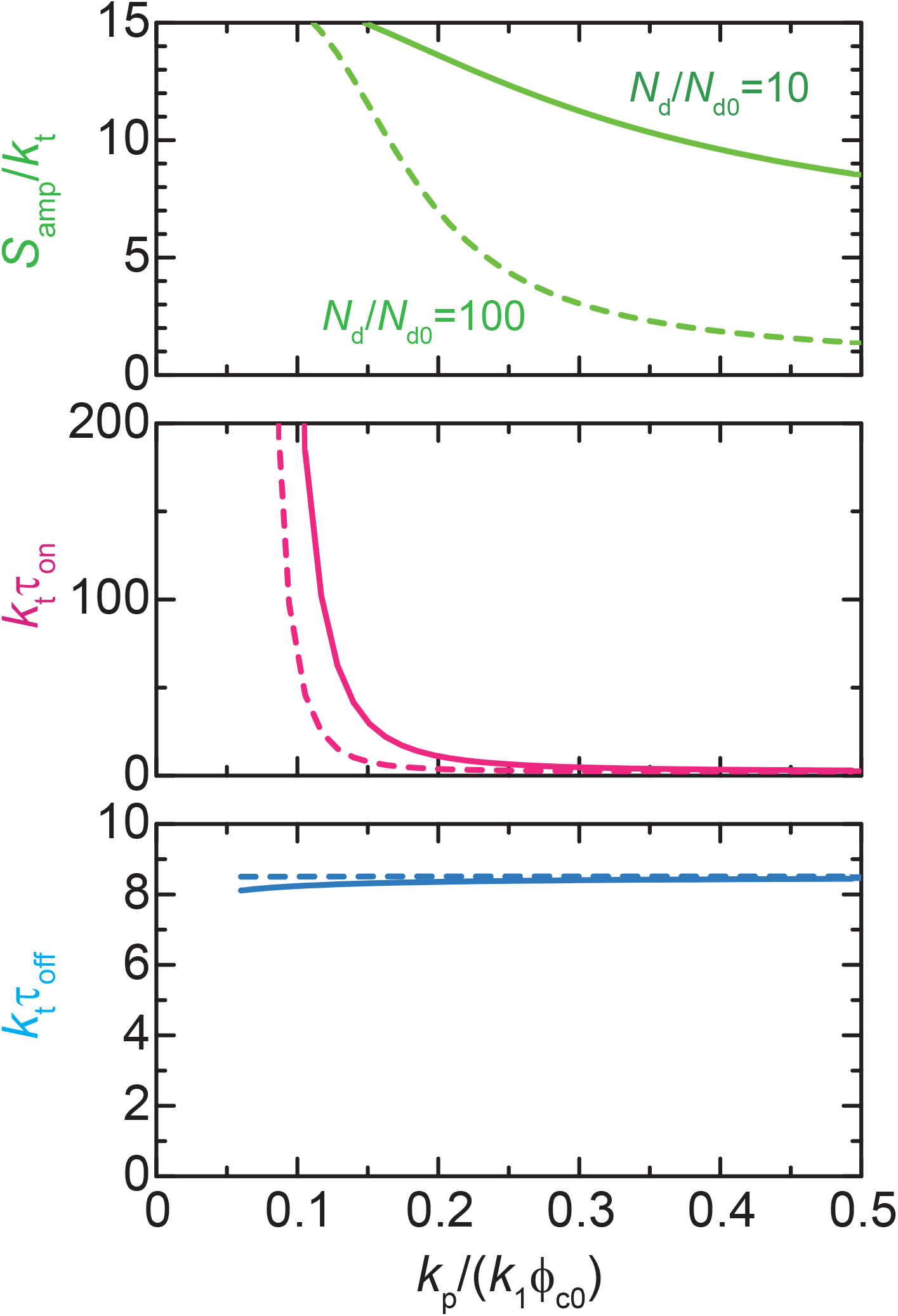
Transcription rate vs phosphorylation rate *k*_p_. The burst size *S*_amp_ (light green), ON duration *τ*_on_ (magenta), and OFF duration *τ*_off_ (cyan) are shown as functions of the rate constant *k*_p_ for the dissociation of complexes from micelle due to the phosphorylation of Pol II C-terminal domains with *N*_d_ = 10.0 (solid) and 100.0 (broken). The values of parameters used for the calculations are summarized in Table 2.

### Pol II-Mediator micelle is stabilized by an enhancer with many binding sites

Using our synthetic enhancer system, we varied the number *n*_e_ of binding sites in the enhancer to quantitatively measure its impacts on the kinetics of transcriptional bursting^18^. In consistent with experimental data, our theory predicts that the transcription rate *S*_st_ at the steady state increases markedly with increasing *n*_e_ (Fig. 6**a**; black solid and dot lines). The burst size and the duration of the ON state increase with increasing *n*_e_ (Fig. 6**b**; green and magenta solid and dot lines, Fig. 6**c**; magenta dot line). It is because the complexes bound to enhancer stabilizes the micelle of Pol II-Mediator complexes. In contrast, the duration of the OFF state decreases with increasing *n*_e_ (Fig. 6**b**; blue solid and dot lines, Fig. 6**c**; blue dot line). This implies that when *n*_e_ is sufficiently large, a stable micelle can be assembled even without additional stabilization from Pol II-Mediator complexes loaded onto the promoter.

**Figure 6.**
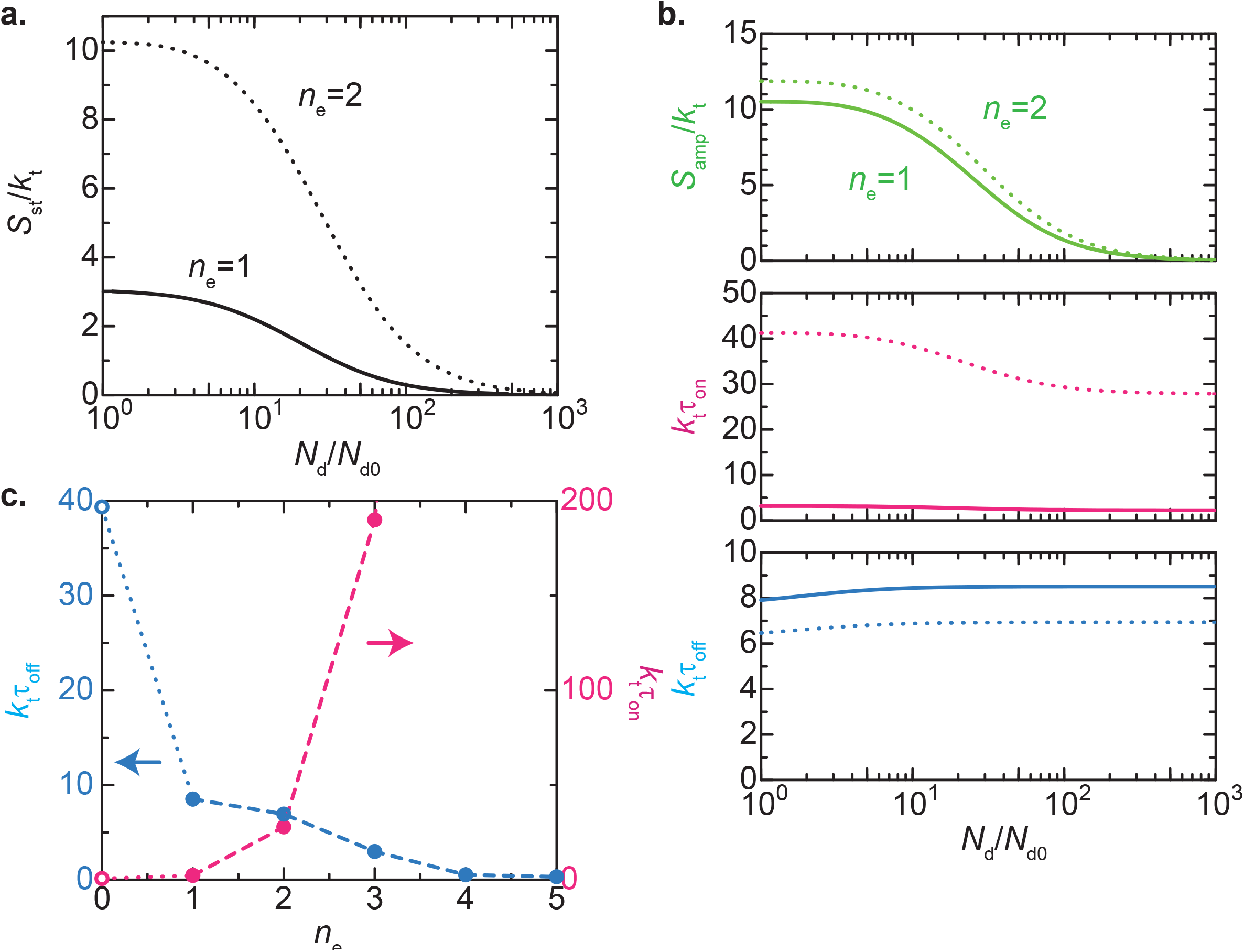
Transcription rate vs number *m*_e_ of enhancer binding sites. **a.** The loading rate *S*_st_ of complexes to the promoter in the steady state (rescaled by rate constant *k*_t_) is shown as a function of the linear enhancer-promoter genomic distance *N*_d_ (rescaled by *N*_d0_) for *n*_e_ = 1 (solid) and 2 (dotted). **b**. The burst size *S*_amp_ (light green), ON duration *τ*_on_ (magenta), and OFF duration *τ*_off_ (cyan) as functions of the linear enhancer-promoter genomic distance *N*_d_ for *n*_e_ = 1 (solid) and 2 (dotted). **c**. The ON duration *τ*_on_ (magenta dots) and OFF duration *τ*_off_ (cyan dots) are shown as functions of the number of enhancer binding sites *n*_e_ for *N*_d_/*N*_d0_ → ∞. The values of parameters used for the calculations are summarized in Table 2.

## Discussion

We have developed a theoretical model that incorporates the connectivity of Pol II-Mediator complexes loaded on to the promoter in the kinetics of transcription reaction as an extension of the theory of polysoap micellization^30^. Transcriptional bursting has been mainly described by using a classical two-state model^46^. Recently, Zuin et al.^14^ and Xiao et al.^47^ applied systems biology approach to explain the non-linear relationship between transcriptional output and enhancer-promoter contact frequency. Both models assume that transcription is enhanced by physical enhancer-promoter contact, either through a regulatory step is triggered upon contact^14^ or through an increasing the number of Pol II molecules in the reaction pool at each contact event^47^. In contrast to these models, we integrate biochemical reaction kinetics with soft matter physics, including the polymeric nature of the chromatin fiber between the enhancer and promoter elements and the stability of micelle assembled by Pol II-Mediator complexes. A key feature of our model is that it quantitatively predicts how these materials properties influence the kinetics of transcriptional bursting, such as burst size and duration. In our framework, the connectivity of Pol II-Mediator complexes bound to enhancers and promoters as well as the cooperativity in micelle assembly and disassembly, rather than enhancer-promoter physical contact, are the primary determinants of transcriptional bursting dynamics.

Our theory identifies the activator clusters observed experimentally as polysoap micelles of Pol II-Mediator complexes. It is motivated by three experimental observations. First, Mediator exhibits adhesive properties due to multivalent Mediator-Mediator interactions^24,25^. Second, Mediator forms complexes by binding to the Pol II CTD^27,28^ and dissociates from Pol II upon phosphorylation of the CTD^39^. Third, Pol II molecules are likely localized at the surfaces of activator clusters to catalyze transcription, based on previous observations that the 3D distance between enhancers and promoters remains relatively large during the ON state^15-17^. Similar implications have been reported for activator clusters formed during zygotic genome activation in zebrafish embryos^48^ and for micro-compartments assembled in the nucleolus^49-51^.

Previously, Mediator has been shown to assemble transcriptional condensates through liquid-liquid phase separation (LLPS)^24,25^. LLPS condensates tend to grow in size to minimize surface energy as long as their components are available, and they remain stable^37,38,40,49^. In contrast, micelles do not grow beyond an optimal size and can be bistable with a disassembled state (Fig. 3**a**) or even unstable^37,38,40,49^. The experimentally observed assembly and disassembly of the activator clusters, along with dynamic transitions between ON and OFF states^18^, may reflect the metastability of micelles. However, this does not exclude the possibilities that LLPS^24,25^ and other types of self-assembly^48-51^ contributes to activator cluster formation. Our theory can be extended to such cases by modifying the free energy (see eq. (1)).

Our theory also offers several experimentally testable predictions. First, both the burst size *S*_amp_ and the duration *τ*_on_ of the ON state increase as the linear genomic distance between enhancer and promoter *N*_d_ decreases (Fig. 4**c**). Second, the kinetic parameters of the transcriptional bursting change at a threshold volume fraction 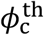 of Pol II-Mediator complexes (Fig. 4**c**). Third, the burst size *S*_amp_ and the ON duration *τ*_on_ increase with decreasing the phosphorylation rate *k*_p_ (Fig. 5) or with increasing the number *n*_e_ of binding sites within the enhancer (Fig. 6**b** and **c**). When *n*_e_ is sufficiently large, micelles assemble at the enhancer without contacting with the promoter. The enhancement of burst size *S*_st_ and duration *τ*_on_ is most pronounced for enhancers with relatively small number *n*_e_. The OFF duration *τ*_off_ is sensitive to the volume fraction *ϕ*_c0_ of complexes (Fig. 4 and Supplementary Figure 3) as well as to the number *n*_e_ and binding energy *ϵ*_d_ of enhancers (Fig. 6**c** and Supplementary Figure 4). It is also expected to depend on material parameters *γ* and *K*, which determine the free energy barrier, although systematic analysis is complicated because of the dependence of the threshold volume fraction 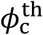 on these parameters. Our theory further predicts the mean-square 3D distance between enhancers and promoters at both ON and OFF states (Supplementary Figure 5). Our live-imaging system^18^ enables quantitative measurement of several of these parameters, as well as precise determination of burst size and durations. It therefore provides an ideal platform to test the prediction of our model.

The key assumption of our theory is that after the loading of Pol II-Med complexes, it takes a finite delay before the subsequent dissociation of Mediator from chromatin. The CTD of *Drosophila* Pol II, composed of 42 YSPTSPS heptapeptide repeats, contains two serine residues per repeat that are phosphorylated during transcription reaction^52^. This assumption can be tested experimentally by using Pol II phosphorylation biosensors, such as Fab^53^ and Mintbody^54^. Indeed, previous live-imaging study suggested that the typical duration of initiation-to-elongation transition corresponds to a phosphorylation rate *k*_p_ greater than 0.01 s^-153^.

To simplify the model and reduce the number of parameters, we made several assumptions. First, we treat Pol II as existing in only two states, hypo-phosphorylated or hyper-phosphorylated, rather than explicitly modeling each phosphorylation site within the CTD. Second, we assume that Pol II forms stable complexes with Mediator^27,28^, with a sufficiently large free energy such that dissociation does not occur until the Pol II CTD is hyper-phosphorylated^39,55^. Third, we neglect enhancer-independent basal transcription from core promoters^56^. Fourth, we assume that the conformational transitions of the chromatin fiber connecting enhancer and promoter occur solely through thermal fluctuations. We did not take account the loop extrusion activity of SMC complexes^57-59^ and the formation of topologically associated domains (TADs)^59,60^ based on previous studies showing that transcription activity is largely uncoupled from TAD formation in *Drosophila* and other speices^45,61-66^. Despite these assumptions and limitations, we believe that our theory captures the essential features of transcriptional bursting regulated by distal enhancers.

Our present theory predicts that the connectivity between Pol II-Mediator complexes loaded at the promoter and those bound at the enhancer catalyzes the assembly of micelles that drive transcriptional bursting. By incorporating the polymeric nature of chromatin fiber and micellar assembly, our theoretical framework extends the classical model in which transcription is activated by physical enhancer-promoter contacts. We envision that this theory serves as a foundation for providing functional insights into the roles of TADs^63,64^, transcriptional condensates^24,25,55,67-69^, superenhancer^24,67-71^, non-coding RNAs^72-74^ in transcriptional regulation. At present, it remains unclear if transcriptional condensates described in previous studies^24,25,55,67-69^ exhibit biophysical properties similar to those of the Pol II-Mediator micelles proposed in our model. Importantly, it has been reported that stable and unstable transcriptional condensates can coexist with in the same nucleus^75^, implying that they do not simply represent LLPS of Mediator and Pol II in the thermodynamic equilibrium. We suggest that further extensions of our theory will provide key functional insights into the biophysical mechanisms underlying transcription regulation by distal enhancers.

## Methods

### Free energy of micelle

The free energy *F*_*m,n*_ of micelle composed of *m* Pol II-Med complexes, among which *n* complexes are loaded to the promoter, has the form^30^

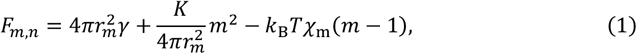

Which is composed of the interfacial energy that represents the multivalent interaction between Mediators that compose the micellar core (first term), the elastic energy that represents the repulsive excluded volume interaction between Pol IIs (second term), the transfer energy of Pol II-Med complexes (third term), and the free energy for the binding between complex and enhancer (fourth term). *γ* is the interfacial energy per unit area and *K* is the elastic constant. *k*_B_*Tχ*_*m*_ is the magnitude of the complex-solvent interaction, relative to the complex-complex interaction. *μ* = 0 at the binodal line of the Pol II-Med complexes (*k*_B_ is the Boltzmann constant and *T* is the absolute temperature). *r*_*m*_ is the radius of the core and has the form

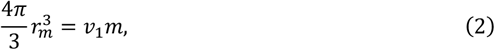

where 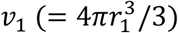 is the volume of Mediator.

### Time evolution equation

We describe the system by using the probabilities, 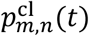 and 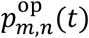, that the micelle is composed of *m* Pol II-Med complexes, among which *n* complexes are loaded on the promoter, at the closed and open DNA conformations, indicated by the superscripts `cl’ and `op’. These probability functions are normalized as

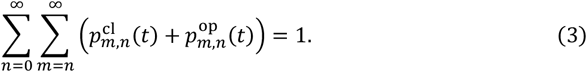

The time evolution equations of 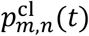 and 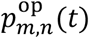 have the forms

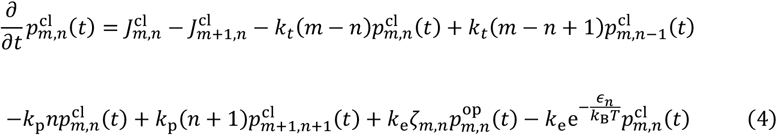

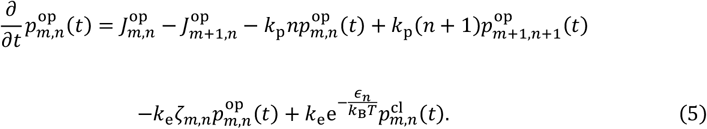

Eq. (4) is composed of the association and dissociation rates of complexes to the micelle by thermal process (first and second terms), the loading rate of complexes to the promoter (third and fourth terms), the dissociation rate of loaded complexes by the phosphorylation of Pol II CTDs (fifth and sixth terms), and the transition of DNA conformation (seventh and eighth terms). Eq. (5) is composed of similar terms, but complexes are not loaded to the promoter because it separates from the micelle at the open DNA conformation. *k*_t_, *k*_p_, and *k*_e_ are the rate constants that account for Pol II loading, Pol II phosphorylation, and DNA conformational transition, respectively.

The fluxes, 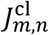 and 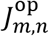, have the forms^33^

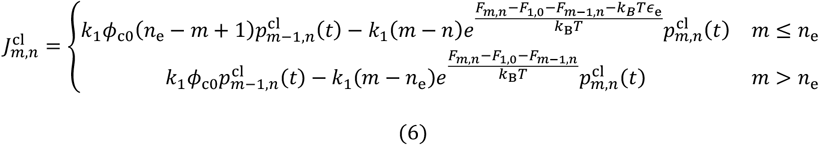

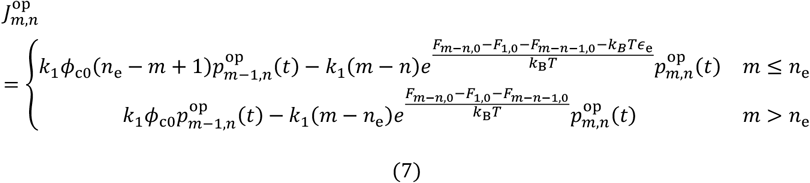

for *n* < *n*_e_ and

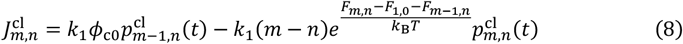

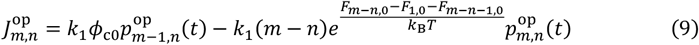

for *n* > *n*_e_. *k*_1_ is the rate constant that accounts for the association and dissociation of complexes. *ϕ*_c0_ is the volume fraction of complexes in the solution.

The probability *ζ*_*m,n*_ that the promoter is at the proximity to the micelle has the form

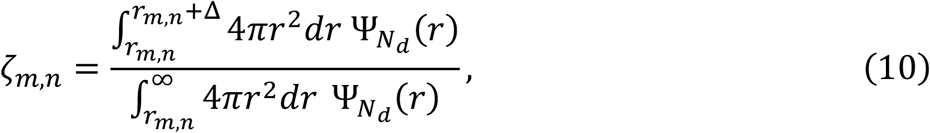

where we used the probability distribution function^34,35,37^

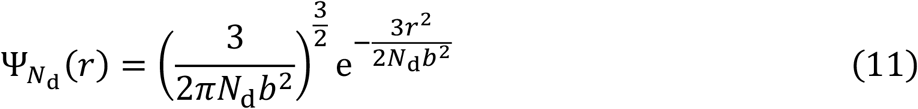

for the Gaussian chain and the minimum distance between the promoter and the micelle,

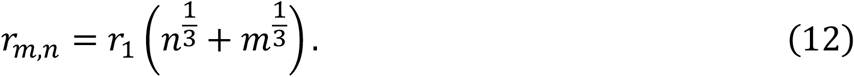

The free energy *ϵ*_n_ necessary to separate the promoter from the micelle has the form

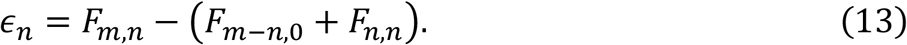

The processes at the longest time scale in eqs. (4) and (5) are the assembly and disassembly of the micelle. Faster processes reach the steady state in the time scales of transcriptional bursting (Supplementary Method).

## Acknowledgement

T.Y. was supported by Grant-in-Aid for Scientific Research (C) (24K06969) and Grant-in-Aid for Transformative Research (A) (20H05934,25H02301) from JSPS (Japan Society for the Promotion of Science). T.F. was supported by the FOREST program (JPMJFR214W) and the CREST program (JPMJCR25T2) from JST (Japan Science and Technology Agency), a Grant-in-Aid for Scientific Research (A) (25H00967), a Grant-in-Aid for Transformative Research Areas (A) (24H02327) from JSPS. K.K. was supported by the JST ACT-X program (JPMJAX2327), a Grant-in-Aid for Transformative Research (A) (25H02441), and a Grant-in-Aid for Early-Career Scientists (24K18090) from JSPS.

## Additional Information

Competing financial interest: The authors declare no financial interest.

## Data availability

The data obtained by the numerical calculations are available in figshare with the identifier (https://doi.org/10.6084/m9.figshare.32038656).

## Code availability

The Mathematica notebook file used for the numerical calculations is available in figshare with the identifier (https://doi.org/10.6084/m9.figshare.32038656).

