## Supplementary Figures and Methods for "Polymeric mechanism of enhancer-promoter cooperativity in transcriptional bursting"

### Supplementary Information: Polymeric mechanism of enhancer-promoter cooperativity in transcriptional bursting

### Supplementary Methods

#### Polymer model of transcriptional burst

To make Supplementary Methods self-contained, we start from the description of the polymer model of transcriptional burst. We consider a stretch of DNA composed of $N_{\rm d}$ Kuhn units between enhancer and promoter. The enhancer is the binding sites of complexes of RNA polymerase II (Pol II) and Mediator (Med). These complexes form a micelle at the enhancer. When the DNA region between enhancer and promoter forms the closed conformation, complexes in the micelle are loaded to the promoter for transcription. The loaded complexes are tethered to the promoter until the C-terminal domains of Pol IIs are heavily phosphorylated so that mediators are dissociated from Pol IIs. The state of the system is represented by the number $m$ of complexes in the micelle, the number $n$ of complexes loaded to the promoter, and the conformational state $-$ closed or open, of the DNA region.

The dynamics of the system is described by the time evolution equations

$$\begin{aligned}\frac{\partial}{\partial t}p^{\rm cl}_{m,n}(t) &= J^{\rm cl}_{m,n} - J^{\rm cl}_{m+1,n} - k_{\rm t}(m-n)p^{\rm cl}_{m,n}(t) + k_{\rm t}(m-n+1)p^{\rm cl}_{m,n-1}(t)\\ &\quad -k_{\rm p}np^{\rm cl}_{m,n}(t) + k_{\rm p}(n+1)p^{\rm cl}_{m+1,n+1}(t)\\ &\quad +k_{\rm e}\zeta_{m,n}p^{\rm op}_{m,n}(t) - k_{\rm e}{\rm e}^{-\beta\epsilon_{m,n}}p^{\rm cl}_{m,n}(t) \qquad \text{(S1)}\\ \frac{\partial}{\partial t}p^{\rm op}_{m,n}(t) &= J^{\rm op}_{m,n} - J^{\rm op}_{m+1,n} - k_{\rm p}np^{\rm op}_{m,n}(t) + k_{\rm p}(n+1)p^{\rm op}_{m+1,n+1}(t)\\ &\quad -k_{\rm e}\zeta_{m,n}p^{\rm op}_{m,n}(t) + k_{\rm e}{\rm e}^{\beta\epsilon_{m,n}}p^{\rm cl}_{m,n}(t), \qquad \text{(S2)}\end{aligned}$$

where $p^{\rm cl}_{m,n}(t)$ and $p^{\rm op}_{m,n}$ are the probabilities that a micelle composed of $m$ complexes is assembled at the enhancer and $n$ of them are already loaded to the promoter in the closed ('cl') and open ('op') DNA conformation, respectively. These probabilities are normalized as

$$\sum_{n=0}^{\infty}\sum_{m=n}^{\infty}(p^{\rm cl}_{m,n}(t) + p^{\rm op}_{m,n}(t)) = 1 \qquad \text{(S3)}$$

Eq. (S1) is composed of terms that represent the association and dissociation of complexes in the micelle by the thermal process (first and second terms), the loading of complexes in the micelle to the promoter (third and fourth terms), the dissociation of complexes due to the phosphorylation of the Pol II C-terminal domains (fifth and sixth terms), and the closing and opening of the DNA region (seventh and eighth terms). Similary, eq. (S2) is composed of terms that represent the association and dissociation of complexes in the micelle by the thermal process (first and second terms), the dissociation of complexes due to the phosphorylation of the Pol II C-terminal domain (third and fourth terms), and the closing and opening of the DNA region (fifth and sixth terms). $k_\mathrm{t}$, $k_\mathrm{p}$, and $k_\mathrm{e}$ are the kinetic constants that account for the loading of complexes to the promoter, the dissociation of complexes due to the phosphorylation of the Pol II C-terminal domains, and the conformational transition of the DNA region, respectively. $\beta$ $(= 1/(k_\mathrm{B}T))$ is the inverse of the thermal energy.

The fluxes, $J^{\mathrm{cl}}_{m,n}$ and $J^{\mathrm{op}}_{m,n}$, of probability have the forms

$$J^{\mathrm{cl}}_{m,n} = \begin{cases} k_1\phi_{\mathrm{c0}}(n_\mathrm{e}-m+1)p^{\mathrm{cl}}_{m-1,n}(t) - k_1(m-n)\mathrm{e}^{\beta(F_{m,n}-F_{1,0}-F_{m-1,n}-k_\mathrm{B}T\epsilon_\mathrm{e})}p^{\mathrm{cl}}_{m,n}(t) & (m \le n_\mathrm{e}) \\ k_1\phi_{\mathrm{c0}}p^{\mathrm{cl}}_{m-1,n}(t) - k_1(m-n_\mathrm{e})\mathrm{e}^{\beta(F_{m,n}-F_{1,0}-F_{m-1,n})}p^{\mathrm{cl}}_{m,n}(t) & (m > n_\mathrm{e}) \end{cases} \tag{S4}$$

$$J^{\mathrm{op}}_{m,n} = \begin{cases} k_1\phi_{\mathrm{c0}}(n_\mathrm{e}-m+1)p^{\mathrm{op}}_{m-1,n}(t) - k_1(m-n)\mathrm{e}^{\beta(F_{m-n,0}-F_{1,0}-F_{m-n-1,0}-k_\mathrm{B}T\epsilon_\mathrm{e})}p^{\mathrm{op}}_{m,n}(t) & (m \le n_\mathrm{e}) \\ k_1\phi_{\mathrm{c0}}p^{\mathrm{op}}_{m-1,n}(t) - k_1(m-n_\mathrm{e})\mathrm{e}^{\beta(F_{m-n,0}-F_{1,0}-F_{m-n-1,0})}p^{\mathrm{op}}_{m,n}(t) & (m > n_\mathrm{e}) \end{cases} \tag{S5}$$

for $n \le n_\mathrm{e}$ and

$$J^{\mathrm{cl}}_{m,n} = k_1\phi_{\mathrm{c0}}p^{\mathrm{cl}}_{m-1,n}(t) - k_1(m-n)\mathrm{e}^{\beta(F_{m,n}-F_{1,0}-F_{m-1,n})}p^{\mathrm{cl}}_{m,n}(t) \tag{S6}$$

$$J^{\mathrm{op}}_{m,n} = k_1\phi_{\mathrm{c0}}p^{\mathrm{op}}_{m-1,n}(t) - k_1(m-n)\mathrm{e}^{\beta(F_{m-n,0}-F_{1,0}-F_{m-n-1,0})}p^{\mathrm{op}}_{m,n}(t) \tag{S7}$$

for $n > n_\mathrm{e}$. Eqs. (S4) - (S7) are composed of the association (first term) and dissociation (second term) rates of complexes. $k_1$ is the rate constant that accounts for the association and dissociation of complexes to the micelle. $\phi_{c0}$ is the volume fraction of complexes in the nucleus. $n_e$ is the number of binding sites in the enhancer.

The dissociation rate is governed by the free energy $F_{m,n}$ of the micelle composed of $m$ complexes, where $n$ complexes are loaded to the promoter. The free energy has the form

$$F_{m,n} = 4\pi r_m^2 \gamma + \frac{K}{4\pi r_m^2} m^2 - k_B T \chi_m (m-1). \tag{S8}$$

Eq. (S8) is composed of the interfacial energy due to the Med-Med multivalent interaction (first term), the Pol II-Pol II excluded volume interaction (second term), and so-called transfer energy, which corresponds to the free energy cost by the interaction between Md and solvent (third term). $\gamma$ is the interfacial energy per unit area, where it accounts for the Med-Med multivalent interaction. $K$ is the elastic constant that accounts for the Pol II-Pol II excluded volume interaction. $k_B T \chi_m$ is the transfer energy. $r_m$ is the radius of the micelle and has the form

$$\frac{4\pi r_m^3}{3} = v_1 m \tag{S9}$$

by using the volume of Mediator $v_1$ $(= 4\pi r_1^3/3)$.

The closing rate of the DNA region is proportional to the probability with which the promoter is located at the proximity to the surface of the micelle

$$\zeta_{m,n} = \frac{\int_{r_{m,n}}^{r_{m,n}+\Delta} 4\pi r^2 dr\, \Psi_{N_d}(r)}{\int_{r_{m,n}}^{\infty} 4\pi r^2 dr\, \Psi_{N_d}(r)}. \tag{S10}$$

where $\Psi_{N_d}(r)$ is the probability distribution function

$$\Psi_{N_d}(r) = \left(\frac{3}{2\pi N_d b^2}\right)^{3/2} e^{-\frac{3r^2}{2N_d b^2}} \tag{S11}$$

of Gaussian chain composed of $N_{\rm d}$ Kuhn units of length $b$. The distance $r_{m,n}$ has the form

$$r_{m,n} = r_1(m^{1/3} + n^{1/3}) \tag{S12}$$

and $\Delta$ is the cutoff distance.

The opening rate is governed by the free energy cost $\epsilon_{m,n}$ to divide a micelle composed of $m$ complexes to two micelles of $m-n$ and $n$ complexes,

$$\epsilon_{m,n} \;=\; F_{m,n} - (F_{n,n} + F_{m-n,0}). \tag{S13}$$

Eq. (S13) is only the free energy difference between the closed and open states and neglects the energy barrier between these states. Our treatment thus provides the lower bound for the free energy cost of the opening of the DNA region. However, practically, the free energy cost $\epsilon_{m,n}$ is relatively large and the opening of the DNA region for $n \neq 0$ is prohibitive.

#### Detailed balance of association and dissociation of complexes

The free energy difference $\mathcal{F}_{m,n}$ typically has one maximum at $m = m^*$, which is sandwitched by two minima $m = m_1$ ($\approx 0$) and $m_2$ ($> m^*$), see Fig. 3 in the main article. The regions at $m \leq m^*$ and $m > m^*$ correspond to the disassembled and assembled states of micelle, respectively. The assembly and disassembly of micelle are rare events, while the association and dissociation of complexes within the disassembled and assembled states are much faster. In the time scale of transcriptional burst, the local equilibrium approximation with respect to the association and dissociation processes is effective at each of the disassembled and assembled states.

With this approximation, the probabilities, $p^{\mathrm{cl}}_{m,n}$ and $p^{\mathrm{op}}_{m,n}$, are written as

$$p^{\mathrm{cl}}_{m,n}(t) = \begin{cases} \frac{\mathrm{e}^{-\beta\mathcal{F}_{m,n}}}{Z_n^{\mathrm{clD}}} q_n^{\mathrm{clD}}(t) & m \leq m^*(n) \\ \frac{\mathrm{e}^{-\beta\mathcal{F}_{m,n}}}{Z_n^{\mathrm{clA}}} q_n^{\mathrm{clA}}(t) & m > m^*(n) \end{cases} \tag{S14}$$

$$p^{\mathrm{op}}_{m,n}(t) = \begin{cases} \frac{\mathrm{e}^{-\beta\mathcal{F}_{m-n,0}}}{Z^{\mathrm{opD}}} q_n^{\mathrm{opD}}(t) & m \leq m^*(0) \\ \frac{\mathrm{e}^{-\beta\mathcal{F}_{m-n,0}}}{Z^{\mathrm{opA}}} q_n^{\mathrm{opA}}(t) & m > m^*(0) \end{cases}, \tag{S15}$$

where $q_n^{\mathrm{clD}}(t)$, $q_n^{\mathrm{clA}}(t)$, $q_n^{\mathrm{opD}}(t)$, and $q_n^{\mathrm{opA}}(t)$ are the probabilities of states with respect to micelle and DNA conformation and are normalized as

$$\sum_{n=0}^{\infty}(q_n^{\mathrm{clD}}(t) + q_n^{\mathrm{clA}}(t) + q_n^{\mathrm{opD}}(t) + q_n^{\mathrm{opA}}(t)) = 1. \tag{S16}$$

The superscripts 'clD'. 'clA', 'opD', and 'opA' stand for the disassembled closed, assembled closed, disassembled open, and assembled open states, respectively. $Z_n^{\mathrm{clD}}$, $Z_n^{\mathrm{clA}}$, $Z^{\mathrm{opD}}$, and $Z^{\mathrm{opA}}$ are the work functions for the corresponding states,

$$Z_n^{\mathrm{clD}} = \sum_{m=n}^{m^*(n)} \mathrm{e}^{-\beta\mathcal{F}_{m,n}} \tag{S17}$$

$$Z_n^{\mathrm{clA}} = \sum_{m=m^*(n)+1}^{\infty} \mathrm{e}^{-\beta\mathcal{F}_{m,n}} \tag{S18}$$

$$Z_n^{\mathrm{opD}} = \sum_{m=n}^{m^*(0)+n} \mathrm{e}^{-\beta\mathcal{F}_{m-n,0}} \tag{S19}$$

$$Z_n^{\mathrm{opA}} = \sum_{m=m^*(0)+n+1}^{\infty} \mathrm{e}^{-\beta\mathcal{F}_{m-n,0}}. \tag{S20}$$

The free energy difference $\mathcal{F}_{m,n}$ has the form

$$\mathcal{F}_{m,n} = F_{m,n} - (m-n)F_{1,0} - F_{n,n} - (m-n)k_{\mathrm{B}}T\log\phi_{\mathrm{c0}} + f^{\mathrm{e}}_{m,n}, \tag{S21}$$

where the free energy $f^{\mathrm{e}}_{m,n}$ with respect to the binding of complexes to the enhancer has the form

$$\beta f^{\mathrm{e}}_{m,n} = \begin{cases} \epsilon_{\mathrm{e}}(m-n) + \log \frac{(m-n)!(n_{\mathrm{e}}-m)!}{(n_{\mathrm{e}}-n)!} & m \le n_{\mathrm{e}} \\ \epsilon_{\mathrm{e}}(n_{\mathrm{e}}-n) + \log(m-n_{\mathrm{e}})! & m > n_{\mathrm{e}} \end{cases} \tag{S22}$$

for $n < n_{\mathrm{e}}$ and

$$\beta f^{\mathrm{e}}_{m,n} = \log(m-n)! \tag{S23}$$

for $n > n_{\mathrm{e}}$.

By using the free energy difference $\mathcal{F}_{m,n}$, eqs. (S4) - (S7) are rewritten as

$$J^{\mathrm{cl}}_{m,n} = k_{m,n}\left(p^{\mathrm{cl}}_{m-1,n}(t) - \mathrm{e}^{\beta\mathcal{F}_{m,n}-\beta\mathcal{F}_{m-1,n}} p^{\mathrm{cl}}_{m,n}(t)\right) \tag{S24}$$

$$J^{\mathrm{op}}_{m,n} = k_{m-n,0}\left(p^{\mathrm{op}}_{m-1,n}(t) - \mathrm{e}^{\beta\mathcal{F}_{m-n,0}-\beta\mathcal{F}_{m-n-1,0}} p^{\mathrm{op}}_{m,n}(t)\right), \tag{S25}$$

where

$$k_{m,n} = \begin{cases} k_1\phi_{\mathrm{c}0}(n_{\mathrm{e}}-m+1) & m \le n_{\mathrm{e}} \text{ and } n \le n_{\mathrm{e}} \\ k_1\phi_{\mathrm{c}0} & \text{otherwise} \end{cases}. \tag{S26}$$

Eqs. (S14) and (S15) satisfy $J^{\mathrm{cl}}_{m,n} = J^{\mathrm{op}}_{m,n} = 0$ within each of the regions, $m \le m^*(n)$ and $m > m^*(n)$, see eqs. (S24) and (S25).

At the boundary of the region, $m = m^*(n)+1$, eq. (S24) is written as

$$\begin{aligned} J^{\mathrm{cl}}_{m^*(n)+1,n} &= k_{m^*(n)+1,n}\left(p^{\mathrm{cl}}_{m^*(n),n}(t) - \mathrm{e}^{\beta\mathcal{F}_{m^*(n)+1,n}-\beta\mathcal{F}_{m^*(n),n}} p^{\mathrm{cl}}_{m^*(n)+1,n}(t)\right) \\ &= k_{m^*(n)+1,n}\mathrm{e}^{-\beta\mathcal{F}_{m^*(n),n}}\left(\frac{1}{Z^{\mathrm{clD}}_n}q^{\mathrm{clD}}_n(t) - \frac{1}{Z^{\mathrm{clA}}_n}q^{\mathrm{clA}}_n(t)\right) \\ &= k^{\mathrm{clA}}_n q^{\mathrm{clD}}_n(t) - k^{\mathrm{clD}}_n q^{\mathrm{clA}}_n(t), \end{aligned} \tag{S27}$$

by noting that $m = m^*(n)$ and $m^*(n)+1$ are at the different states. For simplicity, we write the maximum $\mathcal{F}_{m^*(n),n}$ of the free energy difference as

$$\mathcal{F}_n^* = \mathcal{F}_{m^*(n),n}. \tag{S28}$$

The free energy differences for the disassembled and assembled states are

$$\mathcal{F}_n^{\mathrm{clD}} = -k_{\mathrm{B}}T \log Z_n^{\mathrm{clD}} \tag{S29}$$

$$\mathcal{F}_n^{\mathrm{clA}} = -k_{\mathrm{B}}T \log Z_n^{\mathrm{clA}} \tag{S30}$$

respectively. By using eqs. (S28) - (S30), the assembly and disassembly rates of micelle, $k_n^{\mathrm{clA}}$ and $k_n^{\mathrm{clD}}$, are represented as

$$k_n^{\mathrm{clA}} = k_n^* \mathrm{e}^{-\beta(\mathcal{F}_n^* - \mathcal{F}_n^{\mathrm{clD}})} \tag{S31}$$

$$k_n^{\mathrm{clD}} = k_n^* \mathrm{e}^{-\beta(\mathcal{F}_n^* - \mathcal{F}_n^{\mathrm{clA}})} \tag{S32}$$

with

$$k_n^* = k_{m^*(n)+1,n}. \tag{S33}$$

Similarly, at $m = m^*(0) + n + 1$, eq. (S25) is rewritten as

$$J^{\mathrm{op}}_{m^*(0)+n+1,n} = k_n^{\mathrm{opA}} q_n^{\mathrm{clD}}(t) - k_n^{\mathrm{opD}} q_n^{\mathrm{opA}}(t) \tag{S34}$$

with

$$k_n^{\mathrm{opA}} = k_0^* \mathrm{e}^{-\beta(\mathcal{F}_0^* - \mathcal{F}^{\mathrm{opD}})} \tag{S35}$$

$$k_n^{\mathrm{opD}} = k_0^* \mathrm{e}^{-\beta(\mathcal{F}_0^* - \mathcal{F}^{\mathrm{opA}})}. \tag{S36}$$

The maximum free energy $\mathcal{F}_0^*$ $(= \mathcal{F}_{m^*(0),0})$ and the kinetic constant $k_0^*$ $(= k_{m^*(0)+1,0})$ are defined in eqs. (S28) and (S33), respectively. The free energy differences, $\mathcal{F}^{\rm opD}$ and $\mathcal{F}^{\rm opA}$, have the forms

$$\mathcal{F}^{\rm opD} = -k_{\rm B}T \log Z^{\rm opD} \tag{S37}$$

$$\mathcal{F}^{\rm opA} = -k_{\rm B}T \log Z^{\rm opA}. \tag{S38}$$

Now we derive the time evolution equations for the probabilities, $q_n^{\rm clD}(t)$, $q_n^{\rm clA}(t)$, $q_n^{\rm opD}(t)$, and $q_n^{\rm opA}(t)$. Summing up both sides of eq. (S1) with respect to $m$ for $n \le m \le m^*(n)$ leads to the form

$$\begin{aligned}\frac{\partial}{\partial t} q_n^{\rm clD}(t) &= -k_n^{\rm clA} q_n^{\rm clD}(t) + k_n^{\rm clD} q_n^{\rm clA}(t) \\ &\quad -k_{\rm t}(M_n^{\rm clD} - n) q_n^{\rm clD}(t) + k_{\rm t}(M_{n-1}^{\rm clD} - n + 1) q_{n-1}^{\rm clD}(t) \\ &\quad -k_{\rm p} n q_n^{\rm clD}(t) + k_{\rm p}(n+1) q_{n+1}^{\rm clD}(t) \\ &\quad +k_{\rm e} \zeta_n^{\rm D} q_n^{\rm opD}(t) - k_{\rm e} {\rm e}^{\beta \epsilon_n^{\rm D}} q_n^{\rm clD}(t). \end{aligned} \tag{S39}$$

In eq. (S39), we omitted the term

$$k_{\rm p}(n+1) \sum_{m=m^*(n+1)+1}^{m^*(n)+1} p_{m,n}^{\rm cl}(t), \tag{S40}$$

which is small due to the proximity to the free energy maximum. $M_n^{\rm clD}$, $\zeta_n^{\rm D}$, and ${\rm e}^{-\beta\epsilon_n^{\rm D}}$ are the average of $m$, $\zeta_{m,n}$, and ${\rm e}^{-\beta\epsilon_{m,n}}$ with respect to the number $m$ of complexes in the micelle,

$$M_n^{\rm clD} = \sum_{m=n}^{m^*(n)} m \frac{{\rm e}^{-\beta\mathcal{F}_{m,n}}}{Z_n^{\rm clD}} \tag{S41}$$

$$\zeta_n^{\rm D} = \sum_{m=n}^{m^*(n)} \zeta_{m,n} \frac{{\rm e}^{-\beta\mathcal{F}_{m-n,0}}}{Z^{\rm opD}} \tag{S42}$$

$${\rm e}^{\beta\epsilon_n^{\rm D}} = \sum_{m=n}^{m^*(n)} {\rm e}^{\beta\epsilon_{m,n}} \frac{{\rm e}^{-\beta\mathcal{F}_{m,n}}}{Z_n^{\rm clD}}. \tag{S43}$$

Similarly, the time evolution equations of $q_n^{\mathrm{clA}}(t)$, $q_n^{\mathrm{opD}}(t)$, and $q_n^{\mathrm{opA}}(t)$ are derived as

$$\begin{aligned}
\frac{\partial}{\partial t} q_n^{\mathrm{clA}}(t) &= k_n^{\mathrm{clA}} q_n^{\mathrm{clD}}(t) - k_n^{\mathrm{clD}} q_n^{\mathrm{clA}}(t) \\
&\quad - k_{\mathrm{t}}(M_n^{\mathrm{clA}} - n) q_n^{\mathrm{clA}}(t) + k_{\mathrm{t}}(M_{n-1}^{\mathrm{clA}} - n + 1) q_{n-1}^{\mathrm{clA}}(t) \\
&\quad - k_{\mathrm{p}} n q_n^{\mathrm{clA}}(t) + k_{\mathrm{p}}(n+1) q_{n+1}^{\mathrm{clA}}(t) \\
&\quad + k_{\mathrm{e}} \zeta_n^{\mathrm{T}} q_n^{\mathrm{opD}}(t) + k_{\mathrm{e}} \zeta_n^{\mathrm{A}} q_n^{\mathrm{opA}}(t) - k_{\mathrm{e}} \left( \mathrm{e}^{\beta \epsilon_n^{\mathrm{T}}} + \mathrm{e}^{\beta \epsilon_n^{\mathrm{A}}} \right) q_n^{\mathrm{clA}}(t) \qquad \text{(S44)} \\
\frac{\partial}{\partial t} q_n^{\mathrm{opD}}(t) &= -k^{\mathrm{opA}} q_n^{\mathrm{opD}}(t) + k^{\mathrm{opD}} q_n^{\mathrm{opA}}(t) \\
&\quad - k_{\mathrm{p}} n q_n^{\mathrm{opD}}(t) + k_{\mathrm{p}}(n+1) q_{n+1}^{\mathrm{opD}}(t) \\
&\quad - k_{\mathrm{e}}(\zeta_n^{\mathrm{D}} + \zeta_n^{\mathrm{T}}) q_n^{\mathrm{opD}}(t) + k_{\mathrm{e}} \mathrm{e}^{\beta \epsilon_n^{\mathrm{D}}} q_n^{\mathrm{clD}}(t) + k_{\mathrm{e}} \mathrm{e}^{\beta \epsilon_n^{\mathrm{T}}} q_n^{\mathrm{clA}}(t) \qquad \text{(S45)} \\
\frac{\partial}{\partial t} q_n^{\mathrm{opA}}(t) &= k^{\mathrm{opA}} q_n^{\mathrm{opD}}(t) - k^{\mathrm{opD}} q_n^{\mathrm{opA}}(t) \\
&\quad - k_{\mathrm{p}} n q_n^{\mathrm{opA}}(t) + k_{\mathrm{p}}(n+1) p_{n+1}^{\mathrm{opA}}(t) \\
&\quad - k_{\mathrm{e}} \zeta_n^{\mathrm{A}} q_n^{\mathrm{opA}}(t) + k_{\mathrm{e}} \mathrm{e}^{\beta \epsilon_n^{\mathrm{A}}} q_n^{\mathrm{clA}}(t). \qquad \text{(S46)}
\end{aligned}$$

Eq. (S44) is derived by summing up eq. (S1) with respect to $m$ for $m > m^*(n)$. Eqs. (S45) and (S46) are derived by summing up eq. (S2) with respect to $m$ for $n \le m \le m^*(0) + n$ and $m > m^*(0) + n$, respectively. $M_n^{\mathrm{A}}$, $\zeta_n^{\mathrm{T}}$, $\zeta_n^{\mathrm{A}}$, $\mathrm{e}^{-\beta \epsilon_n^{\mathrm{T}}}$, $\mathrm{e}^{-\beta \epsilon_n^{\mathrm{A}}}$ have the forms

$$M_n^{\mathrm{A}} = \sum_{m=m^*(n)+1}^{\infty} m \frac{\mathrm{e}^{-\beta \mathcal{F}_{m,n}}}{Z_n^{\mathrm{clA}}} \qquad \text{(S47)}$$

$$\zeta_n^{\mathrm{T}} = \sum_{m=m^*(n)+1}^{m^*(0)+n} \zeta_{m,n} \frac{\mathrm{e}^{-\beta \mathcal{F}_{m-n,0}}}{Z^{\mathrm{opD}}} \qquad \text{(S48)}$$

$$\zeta_n^{\mathrm{A}} = \sum_{m=m^*(0)+n+1}^{\infty} \zeta_{m,n} \frac{\mathrm{e}^{-\beta \mathcal{F}_{m-n,0}}}{Z^{\mathrm{opA}}} \qquad \text{(S49)}$$

$$\mathrm{e}^{\beta \epsilon_n^{\mathrm{T}}} = \sum_{m=m^*(n)+1}^{m^*(0)+n} \mathrm{e}^{\beta \epsilon_{m,n}} \frac{\mathrm{e}^{-\beta \mathcal{F}_{m,n}}}{Z_n^{\mathrm{clA}}} \qquad \text{(S50)}$$

$$\mathrm{e}^{\beta \epsilon_n^{\mathrm{A}}} = \sum_{m=m^*(0)+n+1}^{\infty} \mathrm{e}^{\beta \epsilon_{m,n}} \frac{\mathrm{e}^{-\beta \mathcal{F}_{m,n}}}{Z_n^{\mathrm{clA}}}. \qquad \text{(S51)}$$

Eqs. (S1) and (S2) for $p^{\rm cl}_{m,n}(t)$ and $p^{\rm op}_{m,n}(t)$ are thus reduced to eqs. (S39), (S44), (S45), and (S46) for $q^{\rm clD}_n(t)$, $q^{\rm clA}_n(t)$, $q^{\rm opD}_n(t)$, and $q^{\rm opA}_n(t)$.

In the closed DNA conformation, the disassembled state is stable for $0 \le n < n_{\rm sp2}$ and the assembled state is stable for $n > n_{\rm sp1}$. $k^{\rm clD}_n$ and $k^{\rm clA}_n$ are not zero only when both of the states are stable, $n_{\rm sp1} < n < n_{\rm sp2}$. At $n = n_{\rm sp1} - 1$, the time evolution equation of $q^{\rm clD}_n(t)$ has the form

$$\begin{aligned}\frac{\partial}{\partial t} q^{\rm clD}_{n_{\rm sp1}-1}(t) &= -k_{\rm t}(M^{\rm clD}_{n_{\rm sp1}-1} - n_{\rm sp1} + 1) q^{\rm clD}_{n_{\rm sp1}-1}(t) + k_{\rm t}(M^{\rm clD}_{n_{\rm sp1}-2} - n_{\rm sp1} + 2) q^{\rm clD}_{n_{\rm sp1}-2}(t) \\ &\quad - k_{\rm p}(n_{\rm sp1} - 1) q^{\rm clD}_{n_{\rm sp1}-1}(t) + k_{\rm p} n_{\rm sp1} q^{\rm clD}_{n_{\rm sp1}}(t) + k_{\rm p} n_{\rm sp1} q^{\rm clA}_{n_{\rm sp1}}(t) \\ &\quad + k_{\rm e} \zeta^{\rm D}_{n_{\rm sp1}-1} q^{\rm opD}_{n_{\rm sp1}-1}(t) - k_{\rm e} {\rm e}^{\beta \epsilon^{\rm D}_{n_{\rm sp1}-1}} q^{\rm clD}_{n_{\rm sp1}-1}(t). \qquad \text{(S52)}\end{aligned}$$

Eq. (S52) is similar to eq. (S39), except for the fifth term, which represents the disassembly of micelle by the dissociation of a complex by the Pol II-CTD phosphorylation. At $n = n_{\rm sp2} + 1$, the time evolution equation of $q^{\rm clA}_n(t)$ has the form

$$\begin{aligned}\frac{\partial}{\partial t} q^{\rm clA}_{n_{\rm sp2}+1}(t) &= -k_{\rm t}(M^{\rm clA}_{n_{\rm sp2}+1} - n_{\rm sp2} - 1) q^{\rm clA}_{n_{\rm sp2}+1}(t) \\ &\quad + k_{\rm t}(M^{\rm clA}_{n_{\rm sp2}} - n_{\rm sp2}) q^{\rm clA}_{n_{\rm sp2}}(t) + k_{\rm t}(M_{n_{\rm sp2}} - n_{\rm sp2}) q^{\rm clD}_{n_{\rm sp2}}(t) \\ &\quad - k_{\rm p}(n_{\rm sp2} + 1) q^{\rm clA}_{n_{\rm sp2}+1}(t) + k_{\rm p}(n_{\rm sp2} + 2) q^{\rm clA}_{n_{\rm sp2}+2}(t) \\ &\quad + k_{\rm e} \zeta^{\rm T}_{n_{\rm sp2}+1} q^{\rm opD}_{n_{\rm sp2}+1}(t) + k_{\rm e} \zeta^{\rm A}_{n_{\rm sp2}+1} q^{\rm opA}_{n_{\rm sp2}+1}(t) \\ &\quad - k_{\rm e} \left( {\rm e}^{\beta \epsilon^{\rm T}_{n_{\rm sp2}+1}} + {\rm e}^{\beta \epsilon^{\rm A}_{n_{\rm sp2}+1}} \right) q^{\rm clA}_{n_{\rm sp2}+1}(t) \qquad \text{(S53)}\end{aligned}$$

Eq. (S53) is similar to eq. (S44), except for the third term, which represents the assembly of micelle by the loading of Pol II.

#### Local steady state approximation

Eq. (S39) is represents the fact that the probability $q^{\rm clD}_n(t)$ changes with the assembly and disassembly of micelle (first and second terms), the loading of complexes to the promoter (third and fourth terms), the dissociation of complexes by phosphorylation (fifth and sixth terms), and the conformational transition (seventh and eigth terms). Among them, the assembly and disassembly of micelle are probably the slowest processes. We assume that the DNA conformation and the number $n$ of loaded Pol II reach locally steady state with respect to $n$ and DNA conformation quickly after the disassembly and assembly of micelle. With this assumption, the probabilities, $q_n^{\rm clD}(t)$, $q_n^{\rm clA}(t)$, $q_n^{\rm opD}(t)$, and $q_n^{\rm opA}(t)$, are written in the forms

$$q_n^{\rm clD}(t) = \alpha_n^{\rm clD} q_{\rm D}(t) \quad \text{(S54)}$$

$$q_n^{\rm opD}(t) = \alpha_n^{\rm opD} q_{\rm D}(t) \quad \text{(S55)}$$

$$q_n^{\rm clA}(t) = \alpha_n^{\rm clA} q_{\rm A}(t) \quad \text{(S56)}$$

$$q_n^{\rm opA}(t) = \alpha_n^{\rm opA} q_{\rm A}(t), \quad \text{(S57)}$$

where the coefficients, $\alpha_n^{\rm clD}$, $\alpha_n^{\rm opD}$, $\alpha_n^{\rm clA}$, and $\alpha_n^{\rm opA}$, are normalized at each of the states as

$$\sum_{n=0}^{n_{\rm sp2}} \alpha_n^{\rm clD} + \sum_{n=0}^{\infty} \alpha_n^{\rm opD} = 1 \quad \text{(S58)}$$

$$\sum_{n=n_{\rm sp1}}^{\infty} \alpha_n^{\rm clA} + \sum_{n=0}^{\infty} \alpha_n^{\rm opA} = 1. \quad \text{(S59)}$$

The probabilities, $q_{\rm D}(t)$ and $q_{\rm A}(t)$, are normalized as

$$q_{\rm D}(t) + q_{\rm A}(t) = 1. \quad \text{(S60)}$$

To make the following discussion simpler, we define the fluxes

$$I_n^{\rm clD} = \eta_n^{\rm clD} q_{\rm D}(t) \quad \text{(S61)}$$

$$I_n^{\rm opD} = \eta_n^{\rm opD} q_{\rm D}(t) \quad \text{(S62)}$$

$$I_n^{\rm eD} = \eta_n^{\rm eD} q_{\rm D}(t) \quad \text{(S63)}$$

with the coefficients

$$\eta_n^{\mathrm{clD}} = k_{\mathrm{t}}(M_{n-1}^{\mathrm{clD}} - n + 1)\alpha_{n-1}^{\mathrm{clD}} - k_{\mathrm{p}} n \alpha_n^{\mathrm{clD}} \tag{S64}$$

$$\eta_n^{\mathrm{opD}} = -k_{\mathrm{p}} n \alpha_n^{\mathrm{opD}} \tag{S65}$$

$$\eta_n^{\mathrm{eD}} = k_{\mathrm{e}} \zeta_n^{\mathrm{D}} \alpha_n^{\mathrm{opD}} - k_{\mathrm{e}} \mathrm{e}^{\beta \epsilon_n^{\mathrm{D}}} \alpha_n^{\mathrm{clD}}. \tag{S66}$$

By using eqs. (S61) - (S63), eqs. (S39) and (S45) are rewritten as

$$\frac{\partial}{\partial t} q_n^{\mathrm{clD}}(t) = -k_n^{\mathrm{clA}} q_n^{\mathrm{clD}}(t) + k_n^{\mathrm{clD}} q_n^{\mathrm{clA}}(t) + I_n^{\mathrm{clD}} - I_{n+1}^{\mathrm{clD}} + I_n^{\mathrm{eD}} \tag{S67}$$

$$\frac{\partial}{\partial t} q_n^{\mathrm{opD}}(t) = -k^{\mathrm{opA}} q_n^{\mathrm{opD}}(t) + k^{\mathrm{opD}} q_n^{\mathrm{opA}}(t) - k_{\mathrm{e}} \zeta_n^{\mathrm{T}} q_n^{\mathrm{opD}}(t) + k_{\mathrm{e}} \mathrm{e}^{\beta \epsilon_n^{\mathrm{T}}} q_n^{\mathrm{clA}}(t) + I_n^{\mathrm{opD}} - I_{n+1}^{\mathrm{opD}} - I_n^{\mathrm{eD}}. \tag{S68}$$

The local steady state approximation with respect to the loading of complexes, Pol II-CTD phosphorylation, and DNA conformational transition reads

$$\eta_n^{\mathrm{clD}} - \eta_{n+1}^{\mathrm{clD}} + \eta_n^{\mathrm{eD}} = 0 \tag{S69}$$

$$\eta_n^{\mathrm{opD}} - \eta_{n+1}^{\mathrm{opD}} - \eta_n^{\mathrm{eD}} = 0 \tag{S70}$$

with the boundary conditions

$$\eta_1^{\mathrm{clD}} = -\eta_1^{\mathrm{opD}} = \eta_0^{\mathrm{eD}} \tag{S71}$$

$$\eta_{n_{\mathrm{sp2}}}^{\mathrm{clD}} = -\eta_{n_{\mathrm{sp2}}}^{\mathrm{opD}} = -\eta_{n_{\mathrm{sp2}}}^{\mathrm{eD}}. \tag{S72}$$

We derive $\alpha_n^{\mathrm{clD}}$ and $\alpha_n^{\mathrm{opD}}$ by numerically solving eqs. (S69) and (S70).

Similarly, we derive $\alpha_n^{\mathrm{clA}}$ and $\alpha_n^{\mathrm{opA}}$ by numerically solving

$$\eta_n^{\mathrm{clA}} - \eta_{n+1}^{\mathrm{clA}} + \eta_n^{\mathrm{eA}} = 0 \tag{S73}$$

$$\eta_n^{\mathrm{opA}} - \eta_{n+1}^{\mathrm{opA}} - \eta_n^{\mathrm{eA}} = 0 \tag{S74}$$

with

$$\eta_n^{\mathrm{clA}} = k_{\mathrm{t}}(M_{n-1}^{\mathrm{clA}} - n + 1)\alpha_{n-1}^{\mathrm{clA}} - k_{\mathrm{p}} n \alpha_n^{\mathrm{clA}} \tag{S75}$$

$$\eta_n^{\mathrm{opA}} = -k_{\mathrm{p}} n \alpha_n^{\mathrm{opA}} \tag{S76}$$

$$\eta_n^{\mathrm{eA}} = k_{\mathrm{e}} \zeta_n^{\mathrm{A}} \alpha_n^{\mathrm{opA}} - k_{\mathrm{e}} \mathrm{e}^{\beta \epsilon_n^{\mathrm{A}}} \alpha_n^{\mathrm{clA}}. \tag{S77}$$

The boundary conditions imposed to the solution of eqs. (S73) and (S74) have the forms

$$\eta_{n_{\mathrm{sp1}}+1}^{\mathrm{clA}} = -\eta_{n_{\mathrm{sp1}}+1}^{\mathrm{opA}} = \eta_{n_{\mathrm{sp1}}}^{\mathrm{eA}} \tag{S78}$$

$$\eta_{n_{\max}}^{\mathrm{clA}} = -\eta_{n_{\max}}^{\mathrm{opA}} = -\eta_{n_{\max}}^{\mathrm{eA}}. \tag{S79}$$

We set the maximum number $n_{\max}$ of loaded complexes to enable the numerical calculations.

#### Time evolution equation for longest time scale

We first sum up each of eqs. (S67) and (S68) with respect to $n$ over their ranges and further sum them up. This leads to the time evolution equation of $q_{\mathrm{D}}(t)$

$$\begin{aligned} \frac{\partial}{\partial t} q_{\mathrm{D}}(t) &= -k_{\mathrm{A}} q_{\mathrm{D}}(t) + k_{\mathrm{D}} q_{\mathrm{A}}(t) \\ &= -(k_{\mathrm{A}} + k_{\mathrm{D}}) \left( q_{\mathrm{D}}(t) - \frac{k_{\mathrm{D}}}{k_{\mathrm{D}} + k_{\mathrm{A}}} \right) \end{aligned} \tag{S80}$$

with

$$k_{\rm A} = \sum_{n=n_{\rm sp1}}^{n_{\rm sp2}} k_n^{\rm clA}\alpha_n^{\rm clD} + \sum_{n=0}^{\infty} k^{\rm opA}\alpha_n^{\rm clD} - k_{\rm e}\sum_{n=0}^{\infty} \zeta_n^{\rm T}\alpha_n^{\rm opD} + k_{\rm t}(M_{n_{\rm sp2}} - n_{\rm sp2})\alpha_{n_{\rm sp2}}^{\rm clD} \quad \text{(S81)}$$

$$k_{\rm D} = \sum_{n=n_{\rm sp1}}^{n_{\rm sp2}} k_n^{\rm clD}\alpha_n^{\rm clA} + \sum_{n=0}^{\infty} k^{\rm opD}\alpha_n^{\rm opA} + \sum_{n=n_{\rm sp1}}^{\infty} k_{\rm e}{\rm e}^{\beta\epsilon_n^{\rm T}}\alpha_n^{\rm clA} + k_{\rm p}n_{\rm sp1}\alpha_{n_{\rm sp1}}^{\rm clA}. \quad \text{(S82)}$$

The fourth terms of eqs. (S81) and (S82) result from the fact that the disassembled and assembled states are stable for $n < n_{\rm sp2}$ and $n > n_{\rm sp1}$, respectively, see also eqs. (S52) and (S53). $k_{\rm A}$ and $k_{\rm D}$ are the assembly and disassembly rates of micelle, respectively.

#### Burst duration and size

In our theory, the ON- and OFF-states of transcriptional burst correspond to the assembled and disassembled states of micelle, respectively. The durations of ON- and OFF-states thus have the forms

$$\tau_{\rm on} = \frac{1}{k_{\rm D}} \quad \text{(S83)}$$

$$\tau_{\rm off} = \frac{1}{k_{\rm A}}. \quad \text{(S84)}$$

The burst size $S_{\rm amp}$ is estimated by the loading rate of complexes at the ON-state,

$$S_{\rm amp} = k_{\rm t}\sum_{n=n_{\rm sp1}}^{\infty} (M_n^{\rm A} - n)\alpha_n^{\rm clA}. \quad \text{(S85)}$$

In the steady state, $t \to \infty$, the probabilities, $q_{\rm D}(t)$ and $q_{\rm A}(t)$, have the forms

$$q_{\rm D}^{\rm st} = \frac{k_{\rm D}}{k_{\rm D} + k_{\rm A}} = \frac{\tau_{\rm off}}{\tau_{\rm on} + \tau_{\rm off}} \quad \text{(S86)}$$

$$q_{\rm A}^{\rm st} = \frac{k_{\rm A}}{k_{\rm D} + k_{\rm A}} = \frac{\tau_{\rm on}}{\tau_{\rm on} + \tau_{\rm off}}. \quad \text{(S87)}$$

The transcription rate $S_{\mathrm{st}}$ at the steady state is estimated by

$$S_{\mathrm{st}} = k_{\mathrm{t}} \sum_{n=n_{\mathrm{sp1}}}^{\infty} (M_n^{\mathrm{A}} - n)\alpha_n^{\mathrm{clA}} q_{\mathrm{A}}^{\mathrm{st}} + k_{\mathrm{t}} \sum_{n=0}^{n_{\mathrm{sp2}}} (M_n^{\mathrm{D}} - n)\alpha_n^{\mathrm{clD}} q_{\mathrm{D}}^{\mathrm{st}}. \tag{S88}$$

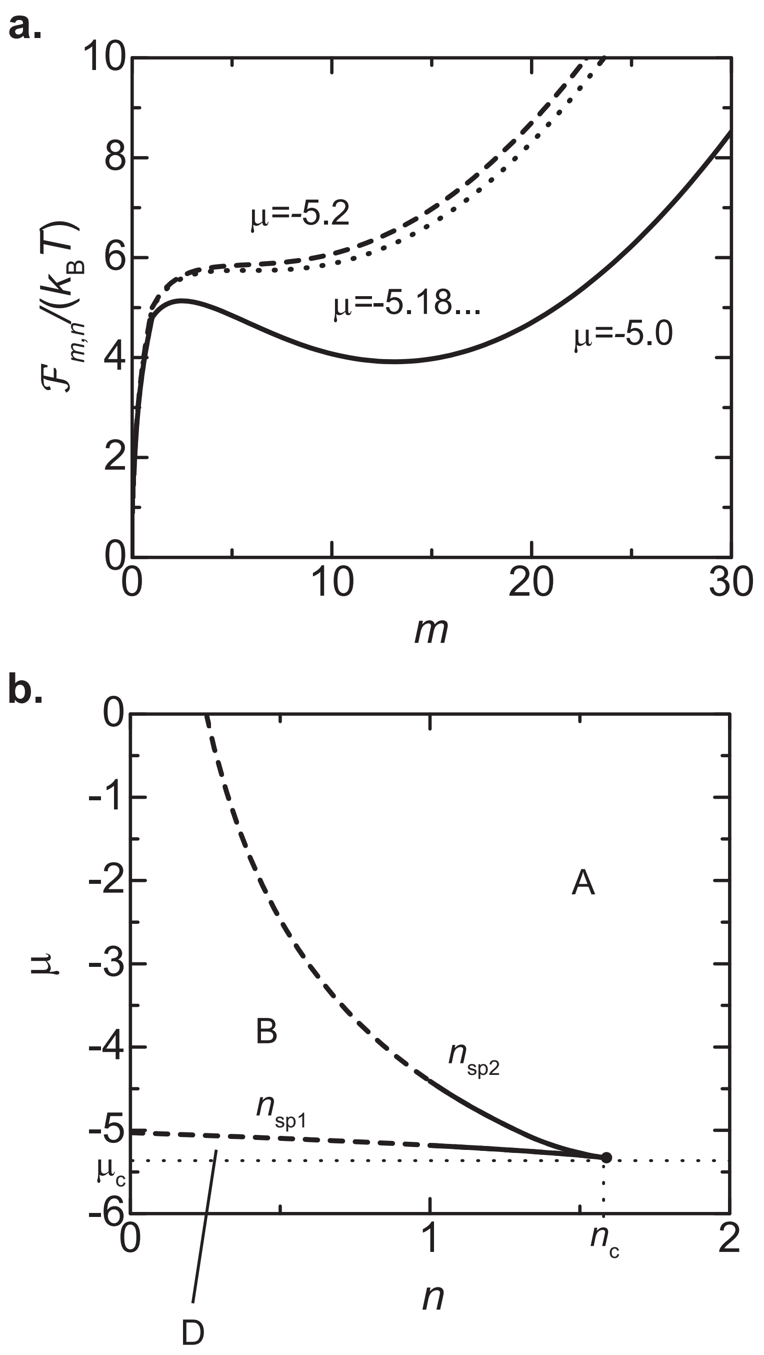


**Supplementary Figure 1 Stability condition of micelle. a.** The free energy difference $\mathcal{F}_{m,n}$ is shown as a function of the number $m$ of Pol II-Med complexes in micelle for $\mu = -5.2$ (solid), $-5.18045$ (dotted), and $-5.2$ (broken). **b.** The stability diagram of micelle with respect to the chemical potential $\mu$ and the number $n$ of Pol II-Med complexes loaded to the promoter. The disassembled state is stable and the assembled state is unstable for $n < n_{\rm sp1}$ (region 'D'). Both of the assembled and disassembled states are stable for $n_{\rm sp1} < n < n_{\rm sp2}$ (region 'B'). The assembled state is stable and the disassembled state is unstable for $n > n_{\rm sp2}$ (region 'A'). The values of parameters used for the calculations are shown in Table 2 in the main article.

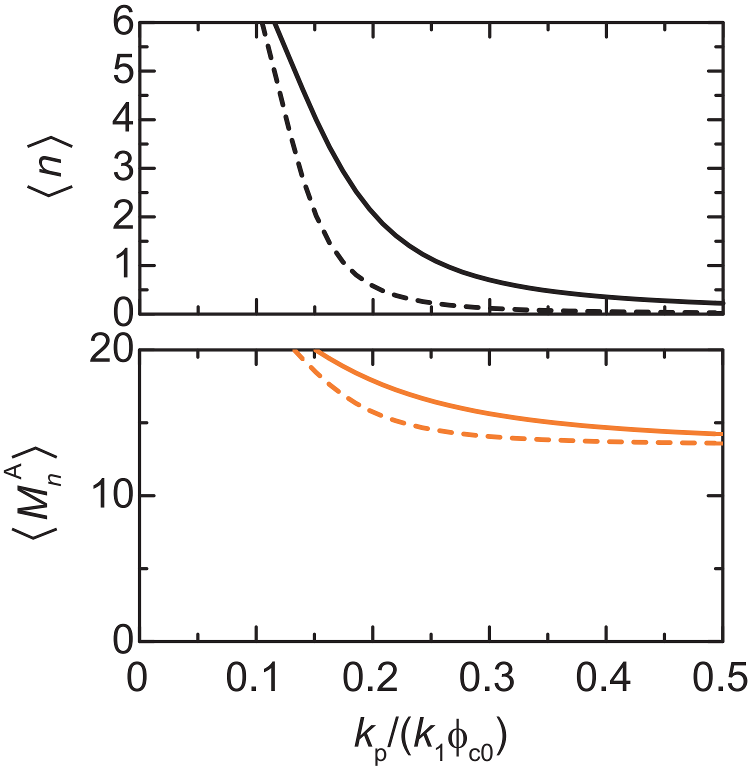


**Supplementary Figure 2 Enhancer-promoter cooperativity.** The mean number $\langle n \rangle$ of Pol II-Med complexes loaded to the promoter (**a**) and the mean number $\langle M_n^{\mathrm{A}} \rangle$ of complexes in the micelle in the assembled state (**b**) are shown as functions of the rate constant $k_{\mathrm{p}}$ that accounts for the phosphorylation of Pol II C-terminal domains (rescaled by $k_1\phi_{\mathrm{c0}}$) for $N_{\mathrm{d}}/N_{\mathrm{d0}} = 10.0$ (solid) and 100.0 (broken). The values of parameters used for the calculations are shown in Table 2 in the main article.

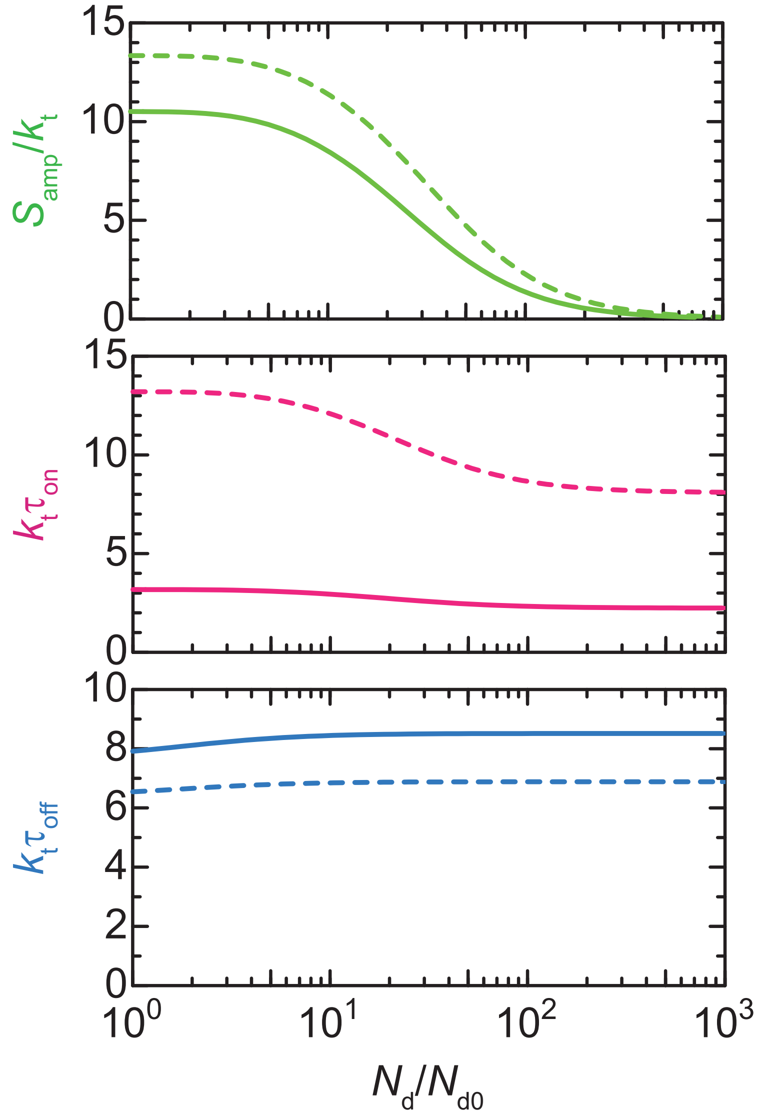


**Supplementary Figure 3 Dependence on chemical potential $\mu$ of Pol II-Med complexes.** The burst size $S_{\mathrm{amp}}$ (light green), ON-duration $\tau_{\mathrm{on}}$ (magenta), and OFF-duration $\tau_{\mathrm{off}}$ (cyan) are shown as functions of the enhancer-promoter genomic distance $N_{\mathrm{d}}/N_{\mathrm{d0}}$ for $\mu = -5.0$ (solid) and $-4.9$ (broken). The values of parameters used for the calculations are shown in Table 2 in the main article.

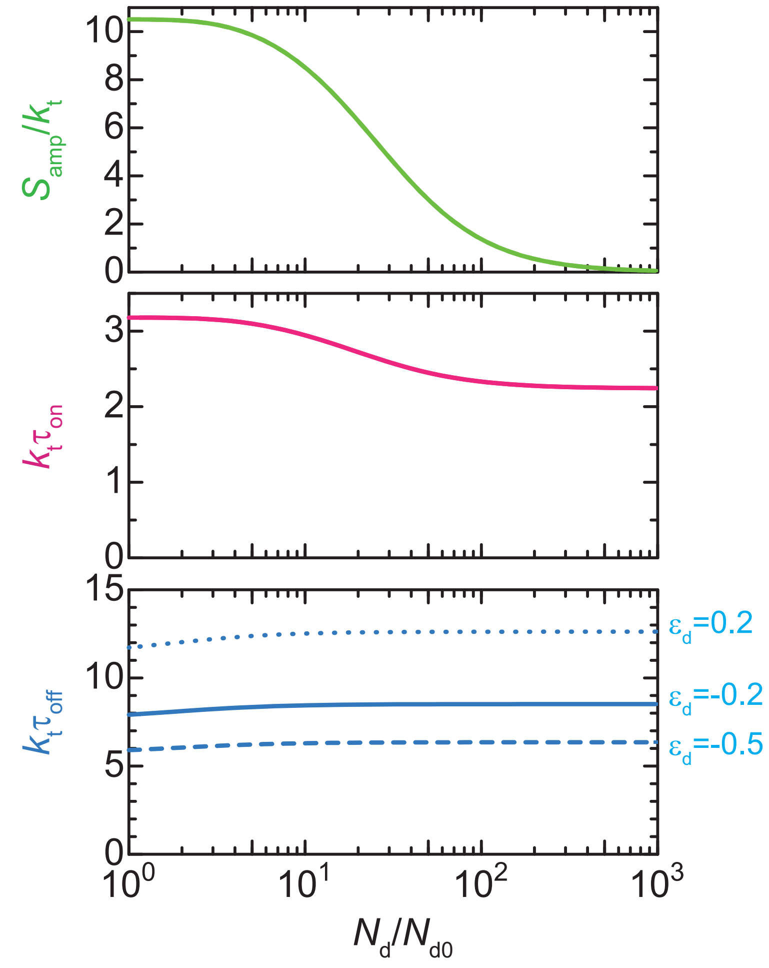


**Supplementary Figure 4 Dependence on enhancer-complex binding energy $\epsilon_d$.** The burst size $S_{amp}$ (light green), ON-duration $\tau_{on}$ (magenta), and OFF-duration $\tau_{off}$ (cyan) are shown as functions of the enhancer-promoter genomic distance $N_d/N_{d0}$ for $\epsilon_d = -0.5$ (broken), $-0.2$ (solid), and 0.2 (dotted). The values of parameters used for the calculations are shown in Table 2 in the main article.

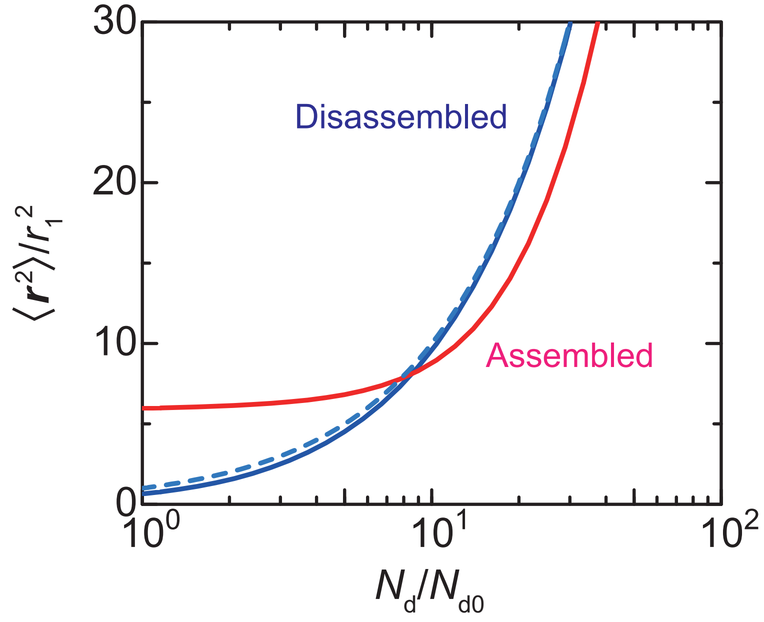


**Supplementary Figure 5 Mean square 3d enhancer-promoter distance vs genomic distance.** The mean square 3d distance $\langle \mathbf{r} \rangle$ between enhancer and promoter is shown as a function of the genomic distance $N_\mathrm{d}$ between these regulatory sequences (rescaled by $N_{\mathrm{d}0}$) for the disassembled (blue solid) and assembled (red solid) states. The blue broken line is the mean square 3d distance for ideal chain. The values of parameters used for the calculations are shown in Table 2 in the main article.
